# Schwann cell plasticity regulates neuroblastic tumor cell differentiation via epidermal growth factor-like protein 8

**DOI:** 10.1101/2020.04.01.019422

**Authors:** Tamara Weiss, Sabine Taschner-Mandl, Andrea Bileck, Fikret Rifatbegovic, Helena Sorger, Max Kauer, Christian Frech, Reinhard Windhager, Christopher Gerner, Peter F. Ambros, Inge M Ambros

## Abstract

The remarkable plasticity of Schwann cells (SCs) enables the acquisition of repair-specific functions essential for peripheral nerve regeneration. We hypothesized that this plastic potential is manifested in stromal SCs found within mostly benign-behaving peripheral neuroblastic tumors. To shed light on the cellular state and impact of stromal SCs, we combined transcriptome and proteome profiling of human ganglioneuromas and neuroblastomas, rich and poor in SC-stroma, respectively, as well as human injured nerve explants, rich in repair SCs. The results revealed a nerve repair-characteristic gene expression signature of stromal SCs. In turn, primary repair SCs had a pro-differentiating and anti-proliferative effect on aggressive neuroblastoma cell lines after direct and trans-well co-culture. Within the pool of secreted stromal/repair SC factors, we identified EGFL8, a matricellular protein with so far undescribed function, to induce neuronal differentiation of neuroblastoma cell lines. This study indicates that human SCs undergo a similar adaptive response in two patho-physiologically distinct situations, peripheral nerve injury and tumor development. This response is mediated by EGFL8 and other SC derived factors, which might be of therapeutic value for neuroblastic tumors and nerve regeneration.

**SYNOPSIS:** 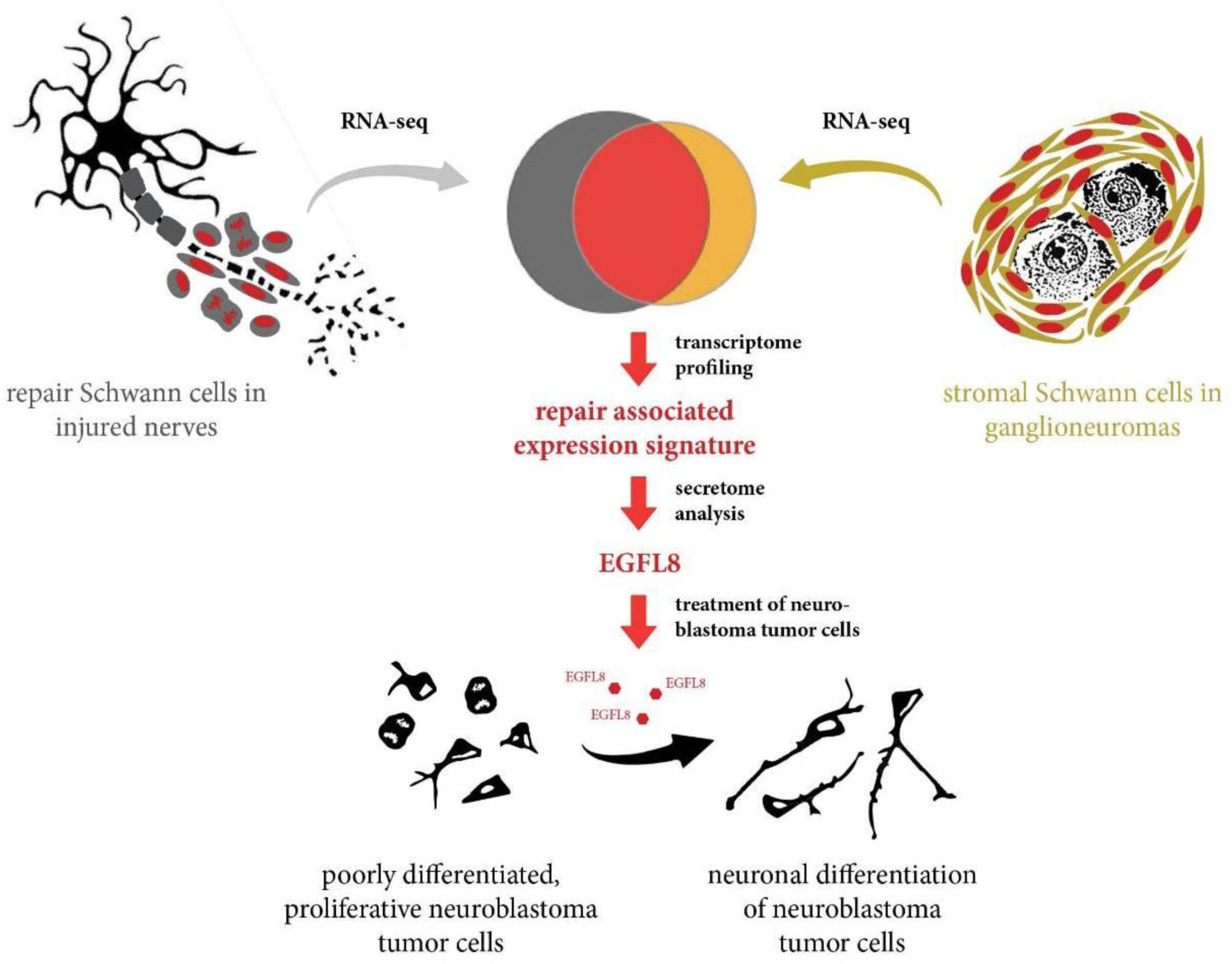

In order to investigate the nature of stromal Schwann cells in benign peripheral neuroblastic tumors (ganglioneuromas), we compared the cellular state of stromal Schwann cells with repair-associated Schwann cells emerging in peripheral nerves after injury.

- Stromal Schwann cells in ganglioneuromas and repair Schwann cells in injured nerves share the expression of nerve repair-associated genes.
- Neuroblastoma cell lines, derived from high-risk metastatic peripheral neuroblastic tumors (neuroblastomas), respond to primary repair Schwann cells and their secretome with increased neuronal differentiation and reduced proliferation.
- Stromal and repair Schwann cells express the matricellular protein EGFL8, which is capable to induce neuronal differentiation of neuroblastoma cell lines in recombinant form.

**THE PAPER EXPLAINED:** *Problem:* In response to peripheral nerve damage, Schwann cells (SCs) are able to transform into specialized repair cells essential for nerve cell regeneration. Our previous studies indicated that this reactive/adaptive potential of human SCs is not restricted to injured nerve cells but also emerges in response to peripheral neuroblastic tumor cells. The usually benign subtypes of peripheral neuroblastic tumors, i.e. ganglioneuroblastomas and ganglioneuromas, contain neuronal differentiating tumor cells and are pervaded by various portions of stromal SCs. Of note, the amount of stromal SCs correlates with a favorable tumor behavior and increased patient survival, whereas aggressive subtypes of peripheral neuroblastic tumors, i.e. neuroblastomas, usually lack stromal SCs and have bad prognosis. This enigma prompted us to investigate the molecular wiring and functional state of stromal SCs *versus* injury-associated repair SCs and how SC signals could be leveraged as therapeutics.

*Result:* Our study revealed that the cellular state of stromal SCs in ganglioneuromas is in many aspects very similar to human repair SCs in injured nerves as both, stromal SCs and repair SCs, are equipped with distinct nerve repair-associated functions. Hence, we exposed different cell lines, derived from high-risk metastatic neuroblastomas, to primary repair SCs or their secretome. The results demonstrated that repair SCs had a pro-differentiating and anti-proliferative effect of on neuroblastoma cell lines upon direct and/or indirect contact. Searching for secreted anti-tumor factors by transcriptome and proteome analyses identified that the matricellular protein EGFL8 was highly expressed in injured nerves and ganglioneuromas. EGFL8 gene expression in peripheral neuroblastic tumors further correlated with increased patient survival. Indeed, treatment of neuroblastoma cell lines with recombinant EGFL8 promoted neuronal differentiation and present EGFL8 as a novel neuritogen.

*Impact:* These findings demonstrate that stromal SCs are equipped with the tools to exert nerve repair-associated functions on peripheral neuroblastic tumor cells and the tumor microenvironment. We further show that the pool of secreted stromal/repair SC molecules contains yet uncharacterized factors with a therapeutic potential for aggressive neuroblastomas. We conclude that the inherent plasticity (reactive/adaptive potential) of SCs is responsible for the development of usually benign ganglioneuroblastomas and ganglioneuromas and, thus, is of utmost interest to be exploited in future treatment approaches for aggressive neuroblastoma subtypes.

## INTRODUCTION

Schwann cells (SCs) are the principal glia of the peripheral nervous system and evolve in close contact with neurons into peripheral nerve fibers. Reciprocal signaling between SCs and neurons regulates the survival, fate decisions, and differentiation of both cell types, but also influences their behavior in regenerative and pathological conditions ^1, 2, 3, 4, 5, 6, 7, 8^. Hence, understanding the molecular mechanisms underlying SC-neuron interaction is of utmost interest to develop effective treatment strategies for injuries and pathologies of the peripheral nervous system.

Despite being necessary for correct nerve development, SCs earned recognition because of their highly plastic cell state that allows them to transform into a dedicated repair cell after peripheral nerve injury. The process is referred to as adaptive cellular reprogramming and enables adult SCs to acquire repair-specific functions, which are essential for nerve regeneration ^9^. This phenotypical switch involves dedifferentiation into a proliferative state with immature/precursor SC properties and redifferentiation along an alternative pathway ^2^. The resulting *repair SC phenotype* represents a transient cell state equipped with a certain skill set specialized to the needs during wound healing such as degradation of myelin debris, attraction of phagocytes, and the expression of cell surface proteins and trophic (neuroprotective and neuritogentic) factors promoting axon re-growth and pathfinding ^2, 10, 11, 12, 13^. Furthermore, we have previously shown that human repair SCs express molecules involved in antigen processing and presentation *via* MHC-II as well as co-inhibitory receptors that support an immunomodulatory role of SCs during nerve regeneration ^13^.

Interestingly, a prevalent stromal SC population is found in usually benign-behaving subtypes of peripheral neuroblastic tumors ^14, 15^. Peripheral neuroblastic tumors originate from trunk neural crest-derived sympathetic neuroblasts ^16, 17^ and are categorized in neuroblastomas (NBs), ganlgioneuroblastomas (GNBs), and ganglioneuromas (GNs) that represent a spectrum from NBs, the most aggressive form, to GNs, the most benign form, and GNBs, which possess various elements of both ^16, 18, 19, 20^. NB and GN subtypes are associated with distinct genomic alterations and strikingly different morphologies ^16, 18^. In general, NBs consist of un- or mostly poorly differentiated tumor cells and cancer-associated fibroblasts ^21^, whereas GNs are composed of differentiated, ganglionic-like tumor cells scattered within a dominant SC stroma ^15, 22^. The content of SC stroma was early recognized as a valuable prognostic factor as it correlates with the degree of tumor cell differentiation and a favorable outcome ^15^. The ganglionic-like tumor cells also extend numerous neuritic processes that form entangled bundles surrounded by ensheathing stromal SCs ^22^. This ganglion-like organoid morphology was assumed to arise from a bi-potent neoplastic neuroblastic precursor cell capable to differentiate along a neuronal and glial lineage ^23^. Hence, an active role of stromal SCs in peripheral neuroblastic tumors has been neglected due to their supposed neoplastic origin.

Of note, previous studies provided evidence for a non-tumor background of stromal SCs ^1, 24^. In a detailed immunohistochemical study, it was shown that the earliest appearance of stromal SCs is confined to the tumor blood vessels and connective tissue septa and not intermingled within the tumor as a clonal origin would imply ^24^. Furthermore, we demonstrated the absence of numerical chromosomal aberrations in stromal SCs, while adjacent ganglionic-like tumor cells possessed a typical aneuploid genome ^1, 25, 26^. These surprising findings argue against the hitherto presumed model of GNB/GN development based on a bi-potent neoplastic cell and support that the tumor cells are able to attract SCs from the nervous environment to the tumor.

In detaching the origin of stromal SCs in GNB/GN from a neoplastic cell, we realized how little we know about their nature. What is the cellular state of stromal SCs? How do they affect GNB/GN development? And why are they not manipulated by the tumor cells to support tumor progression but are associated with a benign tumor behavior/biology? We and others have previously shown that the aggressiveness of NB cell lines, derived from high-risk metastatic NBs, can be reduced upon exposure to SCs and their secreted factors ^27, 28, 29, 30, 31^. However, a comprehensive analysis shedding light on the origin and functional characteristics of stromal SCs is still missing.

Based on the inherent plasticity of SCs and the yet unresolved nature of SC stroma, we speculate that GNB/GN development could be the result of a reactive/adaptive response of SCs to peripheral neuroblastic tumor cells similar to injured nerve cells. Thus, we here compared the cellular state of stromal SCs in GNs to repair SCs in injured nerves by transcriptome profiling of human nerve and human GN tissues. Moreover, we analyzed the effect of human primary repair SCs and their secreted factors on genetically diverse NB cell lines in co-culture studies to identify factors of therapeutic potential for aggressive NBs that lack a SC stroma.

## RESULTS

### Transcriptome profiling revealed that ganglioneuromas contain stromal Schwann cells with a nerve repair-associated gene expression signature and immune cells

To assess the cellular state of stromal SCs, we performed a comprehensive transcriptomic analysis involving human tissues of SC stroma-rich GNs, SC stroma-poor NBs, and repair SC containing injured nerves, alongside with cultures of primary repair SCs and NB cell lines. We confirmed a prevalent SC population in injured nerves (**Fig. 1a**), GNs (**Fig. 1b**) and the absence of SCs in NBs (**Fig. 1c**) by immunofluorescence staining for SC marker S100B. Co-staining with neurofilament heavy polypeptide (NF200) identified degrading axons in injured nerves (**Fig. 1a**) and ganglionic-like tumor cells with abundant neuritic processes in GNs (**Fig. 1b**). According to the un- or poorly-differentiated state of tumor cells in NBs, no NF200 signal was detected in NB samples (**Fig. 1c**). Primary repair SC cultures were positive for S100B and had a purity of >95% (**Fig. 1d**). Cultured NB cell lines highly expressed the neuronal ganglioside GD2 (**Fig. 1e**).

**Figure 1.**
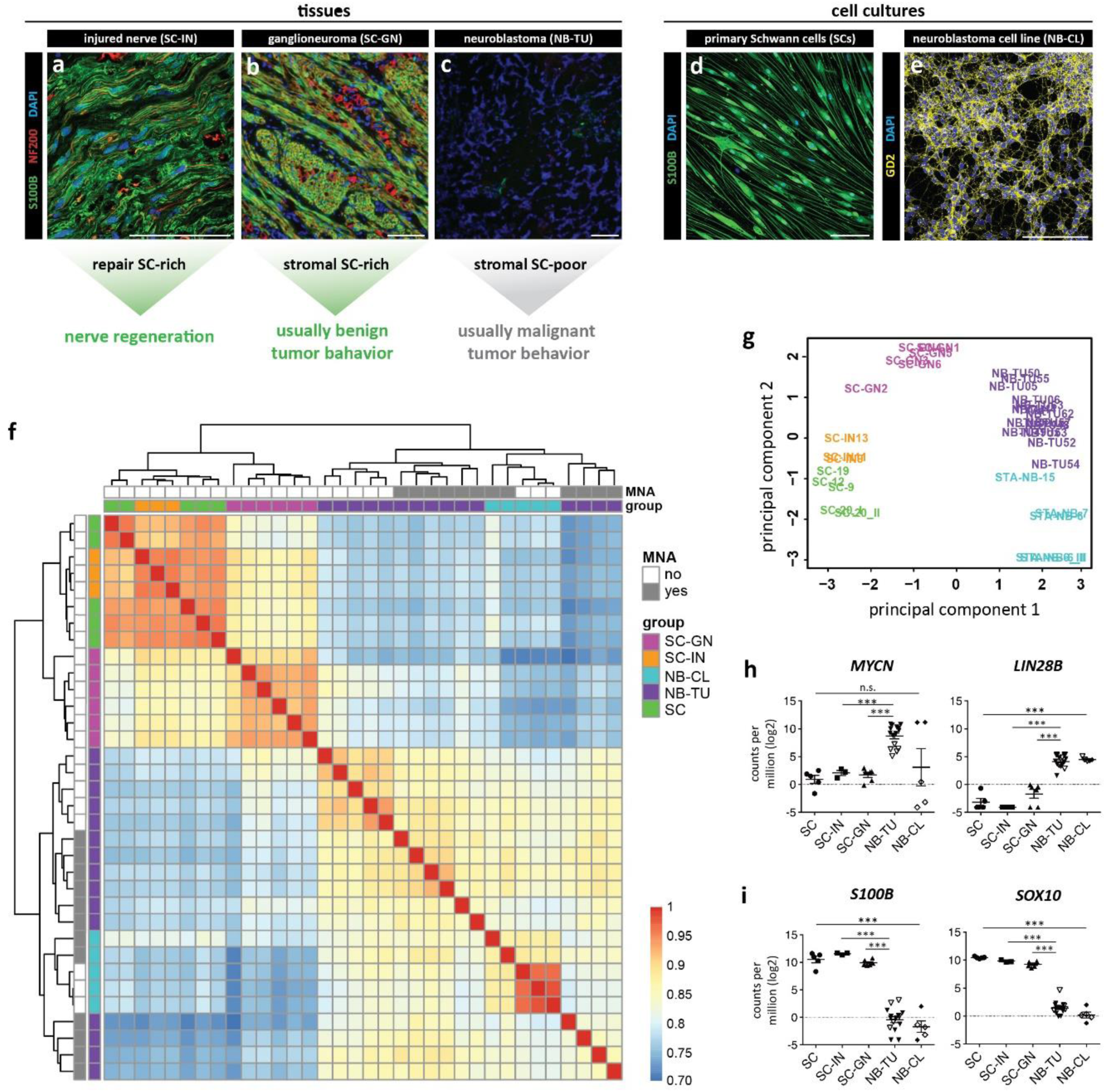
Transcriptome analysis of repair SCs in injured nerves, stromal SCs in ganglioneuromas, neuroblastomas, primary repair SCs, and neuroblastoma cell lines. (a-e) Cell cultures and tissue used for transcriptome profiling; scale bars = 100 µm. Representative cryosections of **(a)** injured nerve fascicle tissue (SC-IN) with S100B positive repair SCs and NF200 positive degrading axons, **(b)** ganglioneuroma tissue (SC-GN) with S100B positive stromal SCs and NF200 positive ganglionic-like tumor cells, and **(c)** neuroblastoma tissue (NB-TU) with NF200 negative tumor cells and no SC stroma. Representative immunofluorescence images of **(d)** human primary Schwann cells (SC) positive for SC marker S100B, and **(e)** the CLB-Ma neuroblastoma cell line (NB-CL) positive for GD2. RNA-seq data of SCs (n=4), NB-CLs (n=3), SC-GN (n=6), SC-INs (n=3) and NB-TU (n=15) illustrated as **(f)** cluster heatmap of sample-to-sample distances and **(g)** principal component analysis (PCA) plot. In the cluster heatmap, sample distances were computed using the Pearson correlation coefficient. Red and blue colors indicate high and low similarity between samples, respectively. Expression level of genes associated with **(h)** aggressive NBs, *MYCN* and *LIN28*, and **(i)** SC lineage genes *S100B* and *SOX10*. Empty symbols indicate *MYCN* non-amplified NB-TUs and NB-CLs. *** q-value ≤ 0.001.

Hierarchical clustering and principal component analysis of obtained RNA-seq data showed that biological samples derived from the same tissue or cell type cluster together and that primary SCs and SC-containing tissues differ from NB cell lines and NB tumors (**Fig. 1f,g**). To confirm tissue/cell identity, we validated the expression of genes associated with either NBs, such as the miRNA suppressor *LIN28B* and the transcription factor *MYCN* ^32, 33^, or the SC lineage, such as *S100B* and *SOX10* ^2^. Indeed, expression of *LIN28B* was significantly higher in NBs and NB cell lines, and the *MYCN* expression level reflected the presence or absence of *MYCN* amplifications in NB cell lines and tumors (**Fig. 1h, S.Table 2&3**). Of note, amplification of the *MYCN* oncogene is associated with an aggressive tumor behavior and poor outcome ^34^. The SC specific genes *S100B* and *SOX10* were significantly and strongly expressed in primary SCs, injured nerves, and GNs (**Fig. 1i**), which is in line with a predominant presence of SCs in these samples and the immunofluorescence results for S100B in Fig. 1a,b&d.

We next defined the characteristic expression signatures of stromal SCs and repair SCs by selecting for genes significantly up-regulated (q-value>0.05; |log2FC|>1) in GNs *versus* NBs, and injured nerves *versus* NBs. In this way, we excluded genes also present in NBs and enriched for genes characteristic for repair SCs in injured nerves and stromal SCs and GNs. Then, we compared the identified expression signatures of stromal SCs and repair SCs, which showed an overlap in 2755 genes (q-value>0.05; |log2FC|>1) (**Fig. 2a**). Functional annotation analysis of these stromal/repair SC genes revealed pathways and gene ontology terms that could be grouped into distinct functional competences. Importantly, these functions reflected the main tasks of human repair SCs in injured nerves involving axon re-growth and pathfinding, lipid/myelin degradation/metabolism, basement membrane formation/ECM (re-)organization, phagocyte attraction, and MHC-II mediated immune regulation ^2, 10, 13, 35^ (**Fig. 2b**). MHC-II expression was validated by immunofluorescence staining of GN sections for HLA-DR. The results showed that HLA-DR was expressed by S100B^+^ stromal SCs, but also indicated the presence of HLA-DR^+^/S100B^-^ immune cells (**S.Fig. 1a-b**). To further analyze a possible repair-related SC state in stromal SCs, we explored the expression of characteristic repair/dedifferentiated SC genes. Indeed, *NGFR, GFAP, ERBB3* and *CADH19* as well as the transcription factors *JUN, ZEB2 and SOX2* ^36, 37, 38^ were significantly up-regulated in SC stroma (**Fig. 2c**). As expected, expression levels of lineage-typical genes were similar in primary cells and corresponding tissue of origin, e.g. primary repair SCs and injured nerves, or NB cell lines and NBs (**Fig. 2b,c**).

**Figure 2.**
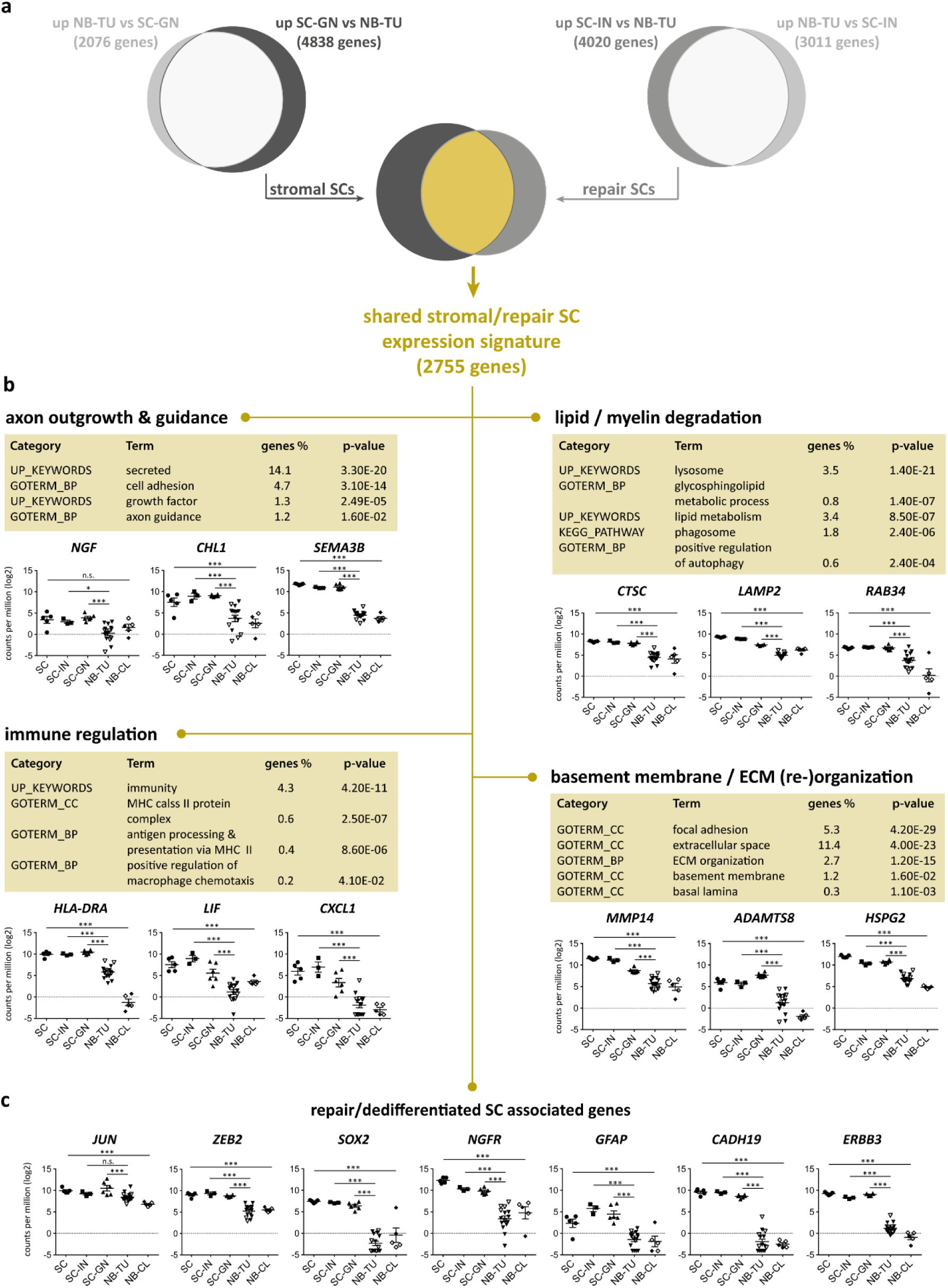
Transcriptome profiling and functional annotation analysis of genes shared by stromal SCs in ganglioneuroma tissue and repair SCs in injured nerve tissue. **(a)** Venn diagrams illustrate the number of significantly regulated genes (q-value>0.05; |log2FC|>1) of stromal SCs (SC-GN) and repair SCs (SC-IN) containing tissues compared to neuroblastoma tissue (NB-TU), respectively, and the overlap in genes shared by stromal and repair SCs (q-value>0.05; |log2FC|>1). **(b)** The DAVID database [37] was used for functional annotation analysis of the 2755 gene set shared by stromal and repair SCs. KEGG pathways, functional categories (UP_KEYWORDS) and gene ontology terms (GOTERM) for biological processes (BP) and cellular compartments (CC) were manually grouped to functions such as axon outgrowth and guidance, lipid/myelin degradation, immune regulation and basement membrane / ECM (re-) organization. The expression of representative genes for each group is shown for all samples. **(c)** Expression of typical repair/dedifferentiated SC associated genes in all samples. Empty symbols indicate *MYCN* non-amplified NB-TUs and NB-CLs. Data are depicted as mean ± SD (n≥3); *** q-value ≤ 0.001, ** q-value ≤ 0.01, * q-value ≤ 0.05, n.s. not significant.

Functional annotation analysis of stromal SC characteristic genes that were not shared with repair SCs revealed an enrichment of gene ontology terms implicated in innate immunity, inflammation as well as T- and B-cell receptor signaling pathways (**S.Table 5**). Accordingly, the presence of T-cells in GN sections was confirmed by the detection of CD3^+^/S100B^-^ T-cells within the CD3^-^/S100B^+^ SC stroma (**S.Fig. 1c-d**). In turn, genes characteristic for repair SCs not shared with stromal SCs were assigned to gene ontology terms for the endoplasmatic reticulum, the Golgi apparatus, vesicle coating and transport, protein transport and binding, as well as acetylation and protein N-linked glycosylation (**S.Table 6**) suggesting an active protein modification and transport machinery.

Taken together transcriptome profiling demonstrated that stromal SCs in GNs and repair SCs in injured nerves share genes associated with distinct nerve repair functions. In addition, we confirmed the presence of CD3^+^ and HLA-DR^+^ immune cells within GNs.

### Direct contact to repair Schwann cells promotes alignment and neurite out-growth of neuroblastoma cells

Since we identified a repair SC-associated gene expression signature in stromal SCs, we used a co-culture model to analyze how NB cells react to repair SCs *in vitro*. Therefore, human primary SC cultures, which reflect all major characteristics of repair SCs ^13^, were isolated, enriched and co-cultured with two human NB cell lines (STA-NB-6 and CLB-Ma) alongside with controls for 11 days (**Fig. 3a**). As a qualitative read-out, we established a multi-color immunofluorescence (IF) staining panel, which identified NB cells by their characteristic expression of the ganglioside GD2, a typical neuronal ganglioside, with low to moderate levels of intermediate filament vimentin (**Fig. 3b**), and SCs by their high expression of both vimentin and SC marker S100B (**Fig. 3c**). After 11 days of co-culture, NB cells had aligned along the bi-polar SC extensions and increased the length of neuritic processes, predominantly in close contact with the SC surface (**Fig. 3d,e** arrows). These results show that the contact to repair SCs induce neuritic out-growth of NB cells *in vitro*.

**Figure 3.**
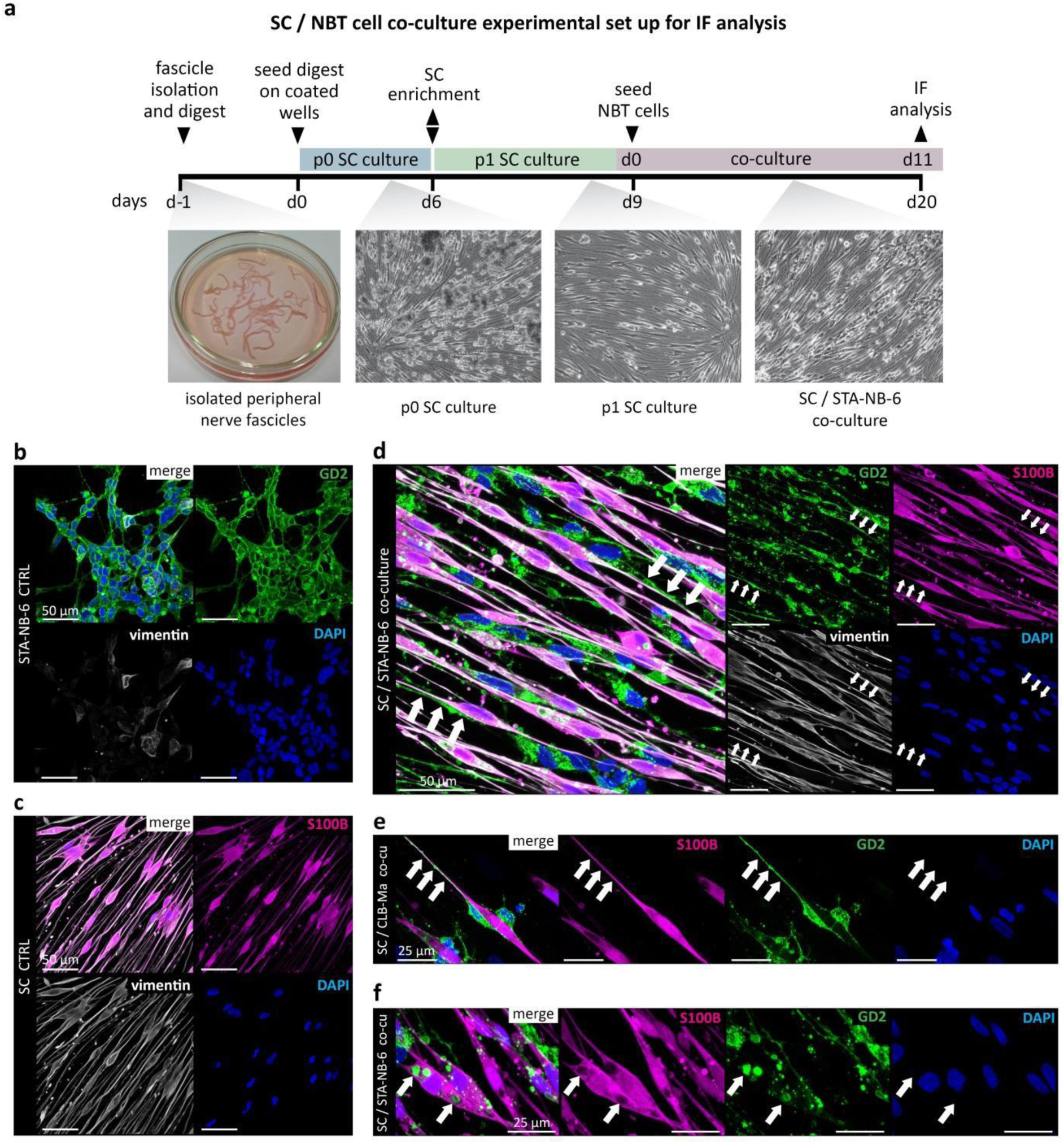
Establishment of a co-culture model to validate the effect of repair SCs on neuroblastoma cell lines *in vitro*. **(a)** Scheme of SC isolation, SC culture, and SC/NB cell co-culture. Representative IF images of **(b)** GD2^+^/vimentin^∼^ STA-NB-6 cells, **(c)** S100B^+^/vimentin^+^ primary repair SCs and **(d-f)** co-cultures with STA-NB-6 or CLB-Ma cells at day 11. Arrows in **(d,e)** indicate extended neuritic processes aligned along SCs. Arrows in **(f)** indicate GD2^+^ droplets localized to S100B^-^ structures within the S100B^+^ cytoplasm of a SC.

In every co-culture we also observed accumulated GD2^+^ signals localized to S100B^-^ vesicular-like structures within the S100B^+^ SC cytoplasm (**Fig. 3f** arrows). As the uptake and degradation of myelin is a key function of repair SCs, the presence of GD2^+^ signals within SCs could indicate their ability to take up lipids different from myelin.

### Repair Schwann cells induce neuronal differentiation of neuroblastoma cells independent of direct cell-cell contact

We next aimed to analyze the effect of primary repair SCs on NB cells using flow cytometry as a quantitative read-out. Therefore, we refined the co-culture settings to distinguish signaling effects between cell bound and secreted molecules. NB cells were either seeded in direct contact with SCs or in a trans-well insert placed above SC cultures allowing diffusion of soluble molecules and reciprocal signaling. The refined co-culture set-up is illustrated in **Fig. 4a**.

**Figure 4.**
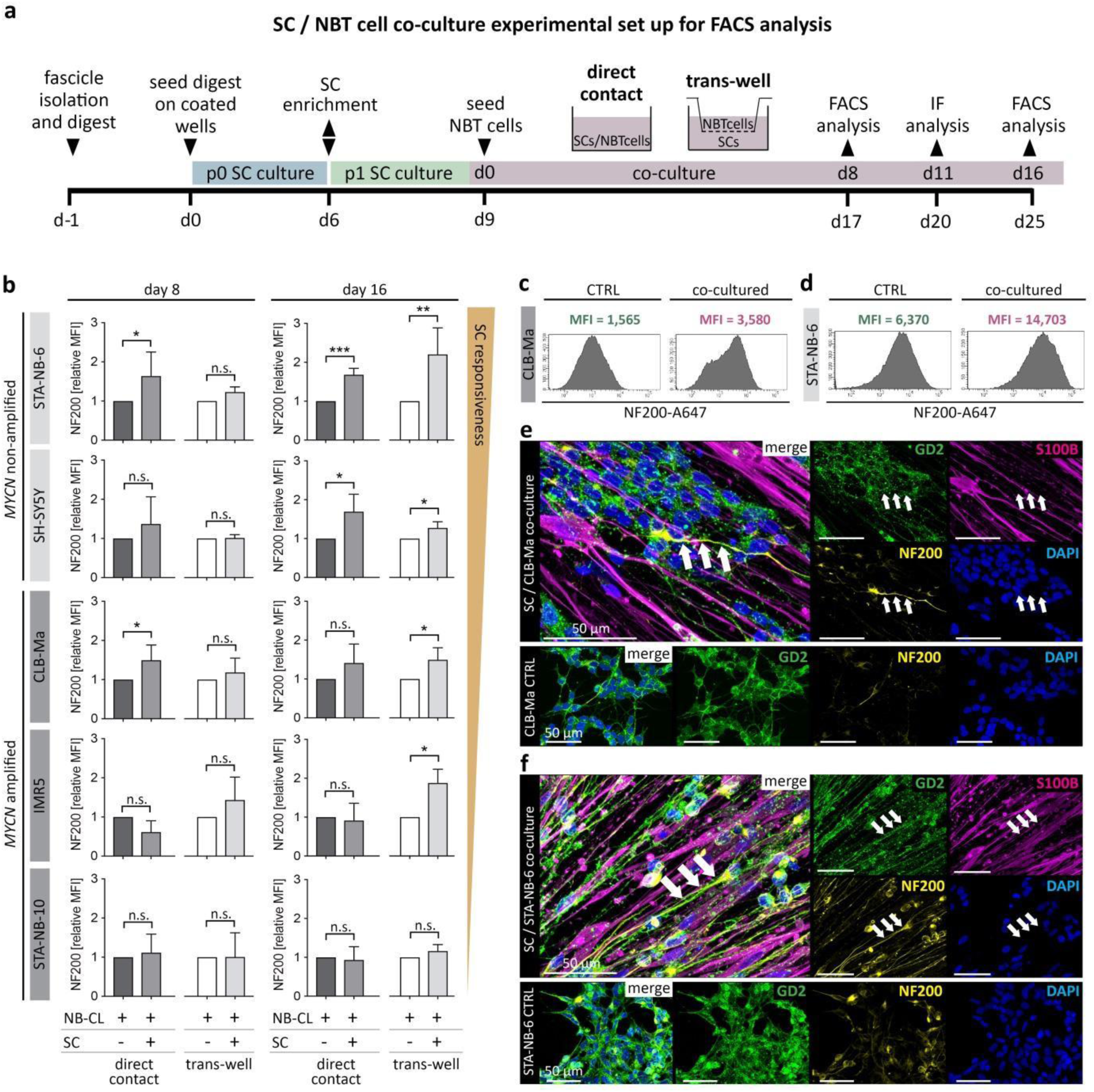
Neuronal differentiation analyses of neuroblastoma cell lines in response to repair SCs *in vitro*. **(a)** Refined SC/NB cell co-culture set up including direct and trans-well co-cultures. Five NB cell lines were co-cultured with primary repair SCs and NF200 expression levels were analyzed by flow cytometry and IF. **(b)** Bar diagrams show the relative mean fluorescence intensity (MFI) of NF200 in GD2^+^S100B^-^ NB cell lines upon direct and trans-well co-culture with SCs at day 8 and day 16. * p ≤ 0.05; ** p ≤ 0.01; *** p ≤ 0.001; n.s. not significant. Representative FACS histograms show the MFI of NF200 in control and co-cultured **(c)** CLB-Ma and **(d)** STA-NB-6 cells at day 16. Representative confocal images of **(e)** CLB-Ma and **(f)** STA-NB-6 cells stained for NF200, S100B, GD2 and DAPI at day 11 of direct co-culture compared to controls. Arrows indicate a NB cell with long neuritic processes strongly positive for NF200.

In order to functionally validate whether repair SCs reenact their key ability of regulating neuronal differentiation on NB cells *in vitro*, five human NB cell lines covering the genetic spectrum of NBs (STA-NB-6, SH-SY5Y, IMR5, STA-NB-10, and CLB-Ma), were co-cultured in direct and indirect contact with human primary repair SCs. After 8 and 16 days, the cultures were analyzed by flow cytometry. The *differentiation FACS panel* discriminated GD2^-^/S100B^+^ SCs and GD2^+^/S100B^-^ NB cells and included the neuronal differentiation marker NF200, which is a major component of the cytoskeleton in mature neurons ^39^. We found that NF200 expression was significantly upregulated in the *MYCN* non-amplified NB cell lines STA-NB-6 and SH-SY5Y after 16 days of direct contact to repair SCs (**Fig. 4b**). Of note, all NB cell lines, except STA-NB-10, showed a significant increase in NF200 expression at day 16 when co-cultured in the trans-wells without direct contact (**Fig. 4b**). The mean fluorescence intensity histograms of NF200 further revealed that the basal NF200 expression level varied among the analyzed cell lines from low, as in CLB-Ma cells (**Fig. 4c, CTRL**), to highest in STA-NB-6 cells (**Fig. 4d, CTRL**). They also demonstrated that the increase in NF200 expression after co-culture was either due to the occurrence of a NF200^+^ subpopulation, e.g. in CLB-Ma cells (**Fig. 4c, co-cultures**), or an overall elevated expression, e.g. in STA-NB-6 cells (**Fig. 4d, co-cultures**). These findings were confirmed by qualitative assessment of NF200 expression by IF stainings of controls and co-cultures of CLB-Ma cells (**Fig. 4e**) and STA-NB-6 cells (**Fig. 4f**).

The results demonstrate that primary repair SCs and/or their secreted factors are sufficient to induce neuronal differentiation of NB cells *in vitro*. We also recognized that the presence or absence of *MYCN* amplification in NB cell lines correlated with their responsiveness to SCs.

### Repair Schwann cells impair proliferation and increase apoptosis of neuroblastoma cells

As cellular differentiation is accompanied by cell cycle arrest, we next determined the proliferation rate of NB cell lines by EdU incorporation in combination with DNA content analysis after direct and trans-well co-culture with SCs. Notably, after 16 days of direct co-culture the number of NB cells in the S-phase was strongly reduced in all tested NB cell lines (**Fig. 5a**). The proliferation rate of trans-well co-cultures was also significantly decreased in all NB cell lines, except STA-NB-10, but less pronounced as upon direct contact (**Fig. 5a**). The strongest anti-proliferative effects were detected in *MYCN* non-amplified cell lines STA-NB-6 and SH-SY5Y as well as *MYCN* amplified IMR5 (**Fig. 5a**). Representative FACS plots illustrated the reduction of proliferation in CLB-Ma cells (**Fig. 5b**) and almost absent proliferation in STA-NB-6 cells (**Fig. 5c**) after 16 days of co-culture. This was validated by IF staining of controls and direct co-cultures of CLB-Ma cells (**Fig. 5d**) and STA-NB-6 cells (**Fig. 5e**) including the proliferation marker Ki67.

**Figure 5.**
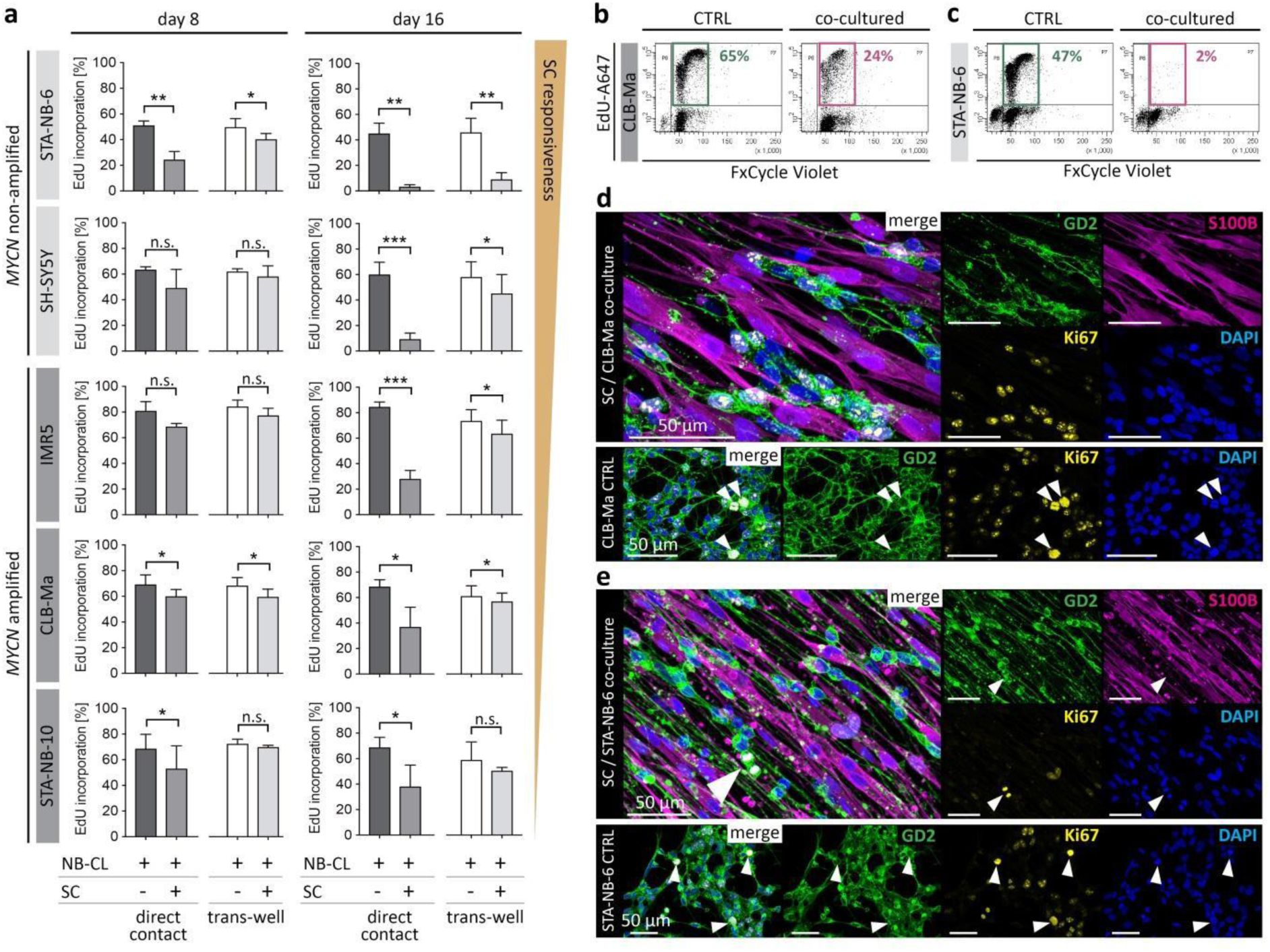
Proliferation analysis of neuroblastoma cell lines after direct and indirect contact to repair SCs *in vitro*. Five NB cell lines were co-cultured with primary repair SCs and their proliferation rates were analyzed by flow cytometry and IF. **(a)** Bar diagrams show the mean percentage of EdU-incorporation ± SD (n≥3) in GD2^+^/S100B^-^ NB cell lines upon direct and trans-well co-culture with SCs at day 8 and day 16; * p ≤ 0.05; ** p ≤ 0.01; *** p ≤ 0.001. Representative FACS plots illustrate EdU incorporation and the DNA content of control and co-cultured NB cell lines **(b)** CLB-Ma and **(c)** STA-NB-6 at day 16; the marked EdU^+^/FxCycleViolet^+^ cells are in the S-Phase of cell cycle. Representative confocal images of **(d)** CLB-Ma and **(e)** STA-NB-6 cells stained for Ki67, S100B, GD2 and DAPI at day 11 of direct co-culture compared to controls; arrows indicate NB cells undergoing mitosis.

In addition to increased differentiation and impaired proliferation, also cell death contributes to the decrease of tumor cells during GN development. Hence, we performed a terminal deoxynucleotidyl transferase dUTP nick end labeling (TUNEL) assay in combination with IF staining for GD2 and S100B to detect apoptotic NB cells in control and co-cultures (**S.Fig. 2a,b**). Quantitative evaluation showed that the apoptosis rate of both, *MYCN* non-amplified STA-NB-6 and *MYCN* amplified CLB-Ma cells, was increased about 10% at day 11 after direct co-culture (**S.Fig. 2c**).

Taken together, these findings show that direct and/or indirect contact to repair SCs decreased the proliferation of NB cell lines. Human repair SCs also elevated the apoptosis rate of NB cell lines upon direct contact. Furthermore, the *MYCN* amplification status correlated with the responsiveness of NB cells to SCs and revealed STA-NB-6 as the strongest and STA-NB-10 as the weakest SC-responsive NB cell line tested.

### Stromal and repair Schwann cells express EGFL8, which is able to induce neurite outgrowth and neuronal differentiation of neuroblastoma cells

After demonstrating a pro-differentiating and anti-proliferative impact of human primary repair SCs on NB cells *in vitro*, we next aimed to identify the factors able to mediate these effects. Therefore, we interrogated the set of transcripts shared by repair SCs in injured nerve tissue and stromal SCs in GN tissue for the expression of secreted factors. Factors of interest were prioritized according to literature research and whether associated receptors, if known, were expressed by NBs. The shared secretome of repair and stromal SCs included neurotrophins such as *NGF, BDNF* and *GDNF* that confirmed the validity of our approach (**Fig. 6a**). In addition, we identified further highly expressed factors of interest such as *IGFBP6, FGF7* and *EGFL8* (**Fig. 6a**). IGFBP-6 was previously reported to inhibit the growth of SH-SY5Y cells ^40^ and FGF7 is involved in neuromuscular junction development ^41^, but both factors were not yet associated with SCs. Notably, EGFL8 was recently described by us as a potential factor involved in nerve regeneration but with yet unknown function ^13^. Other neurotrophic factor transcripts, such as *PTN*, highly expressed in stromal but not in repair SCs, and *CNTF*, expressed in repair but not in stromal SCs, were included in the panel of candidate factors as transcripts of their putative receptors were present in NBs (**S.Fig. 3**).

**Figure 6.**
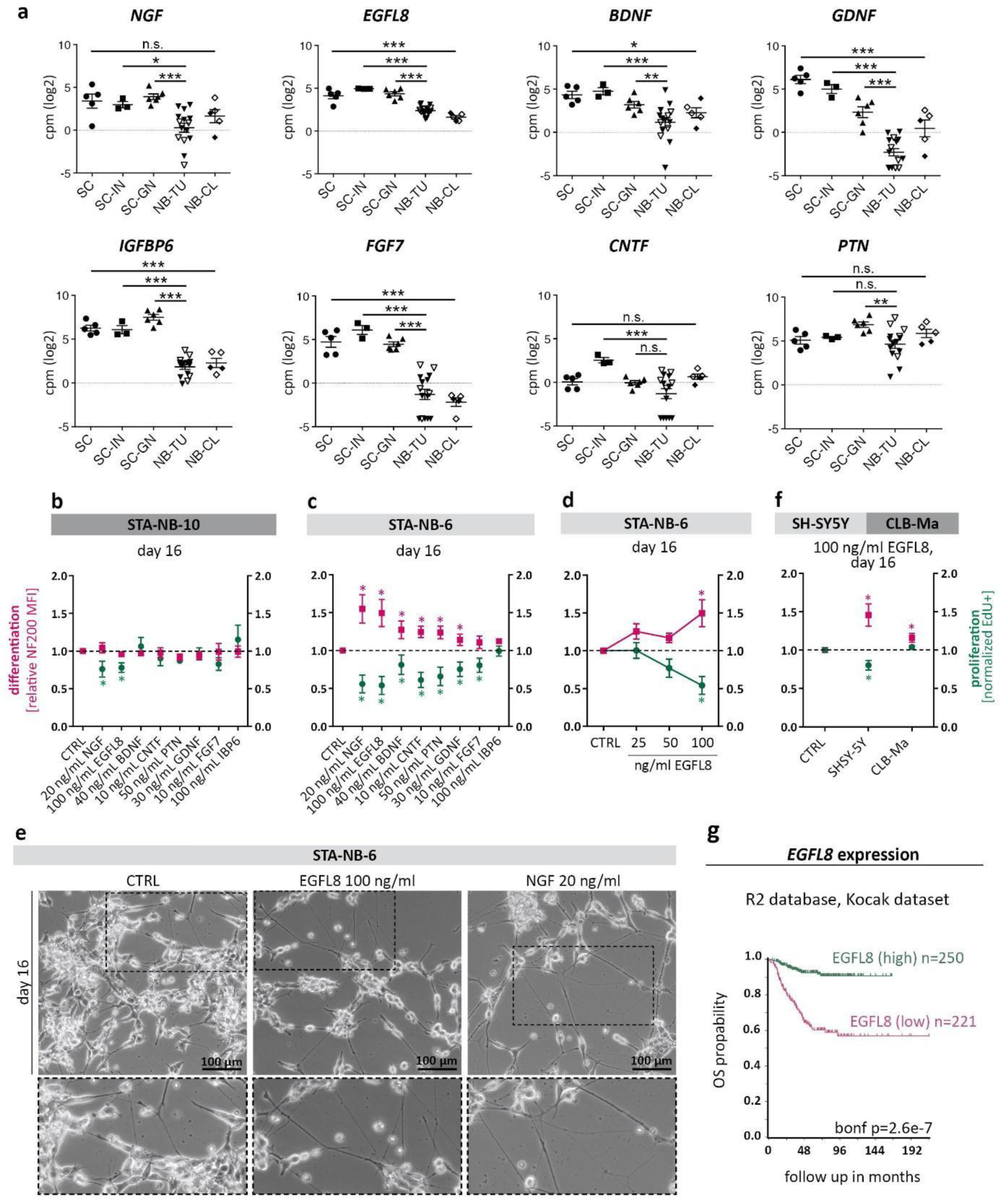
Neuronal differentiation and anti-proliferative effects of secreted factors shared by stromal and repair SCs. **(a)** Expression levels of chosen candidate factors *NGF, EGFL8, BDNF, GDNF, IGFBP6, FGF7, CNTF* and *PTN* in primary repair SCs (SC), injured nerve fascicle tissue (SC-IN), SC stroma rich GN tissue (SC-GN), NB tissue (NB-TU) and NB cell lines (NB-CL); lined symbols indicate *MYCN* non-amplified non-amplified NB-TUs and NB-CLs. ** p ≤ 0.01***, p ≤ 0.001, n.s. not significant. Relative NF200 MFI and normalized EdU incorporation measured by FACS analysis of **(b)** STA-NB-10 and **(c)** STA-NB-6 cells cultured for 16 days in the absence or presence of recombinant proteins at concentrations as indicated. Medium containing recombinant proteins was replenished every 3-4 days. **(d)** STA-NB-6 cells cultured for 16 days in the absence or presence of EGFL8 at concentrations as indicated and analyzed by FACS. **(e)** Bright field images of STA-NB-6 cells at day 16 of culture in the absence or presence of 100 ng/ml EGFL8 or 20 ng/ml NGF, respectively. Enlargements illustrate increased outgrowth of neuritic processes in EGFL8 and NGF treated cultures. **(f)** SH-SY5Y and CLB-Ma cell lines after exposure to 100 ng/ml EGFL8 and untreated cultures. (b, c, d, f) * p ≤ 0.05; n≥3. **(g)** Kaplan-Meier survival plots show the overall survival (OS) probability of patients grouped according to high and low *EGFL8* expression in primary tumors at diagnosis. Data were derived from the Kocak dataset of the *R2 Genomics Analysis and Visualization platform (see also SF2)*.

In order to validate the effect of a set of 8 candidate factors, the recombinant proteins NGF, BDNF, GDNF, CNTF, PTN, FGF7, IBP6 and EGFL8 were added to the SC-weakly-responsive STA-NB-10 and SC-strongly-responsive STA-NB-6 cells. Proliferation and neuronal differentiation were monitored by flow cytometry after 16 days of exposure to respective factors. As expected, the factors had less impact on the SC-weakly-responsive cell line STA-NB-10, however, NGF and EGFL8 caused a significant anti-proliferative effect (**Fig. 6b**). In contrast, the SC-strongly-responsive STA-NB-6 cells were significantly impaired in proliferation and showed increased neuronal differentiation after treatment with either NGF, EGFL8, BDNF, CNTF, PTN or GDNF (**Fig. 6c**). Notably, the effect of EGFL8 was concentration dependent and comparable to NGF, one of the most potent neurotrophins known so far (**Fig. 6d**). Phase contrast images confirmed a reduction of cell number and increase in the length of neuritic processes in NGF-as well as EGFL8-treated STA-NB-6 cells (**Fig. 6e**). EGFL8 also acted pro-differentiating on CLB-Ma and SH-SY5Y cells, while an anti-proliferative effect was only observed in the latter (**Fig. 6f**).

We demonstrate that EGFL8, a protein so far only described in thymocyte development ^42^, represents a novel neuritogenic factor able to enhance neuronal differentiation and/or to impair proliferation of aggressive NB cell lines.

### The *EGFL8* gene expression level in neuroblastomas correlates with increased patient survival

As EGFL8 exerted anti-tumor activity on NB cells *in vitro*, we next assessed whether *EGFL8* expression levels in peripheral neuroblastic tumors may correlate with the clinical outcome. Analysis of the overall patient survival (OS) according to *EGFL8* gene expression was performed using the *R2: Genomics Analysis and Visualization platform*. Two different datasets, comprising 498 and 649 tumor specimens, respectively, demonstrated an over 90% OS probability for patients with high *EGFL8* expression, but it was less than 60% for patients with low *EGFL8* expression (**Fig. 6g & S.Fig. 4**). Unfortunately, these datasets did not provide information about the stromal SC content of the included tumor specimens. It is likely that EGFL8 expression is not exclusive for stromal SCs but its correlation with increased patient survival could be due to its neuritogenic effect on peripheral neuroblastic tumor cells.

### EGFL8 protein abundance is significantly higher in ganglioneuromas than in neuroblastomas

To verify whether the high *EGFL8* gene expression detected in GNs is reflected in the EGFL8 protein level, we performed high-resolution mass spectrometry analysis of SC stroma-rich GNs as well as SC stroma-poor NBs and analyzed this new data set together with our existing proteomic data set comprising NB cell lines, repair SC-containing injured nerve tissue as well as primary repair SC cultures ^13^. The results show a significantly higher abundance of the EGFL8 protein in injured nerves and GNs when compared to NBs (**Fig. 7a**). In addition, the protein levels of EGFL8 in primary cells reflected their respective tissue of origin (**Fig. 7a**). Hence, mass spectrometric analyses confirmed that the EGFL8 protein is highly abundant in SC stroma-rich GNs and repair SC-rich injured nerve tissues as well as primary repair SC cultures.

**Figure 7.**
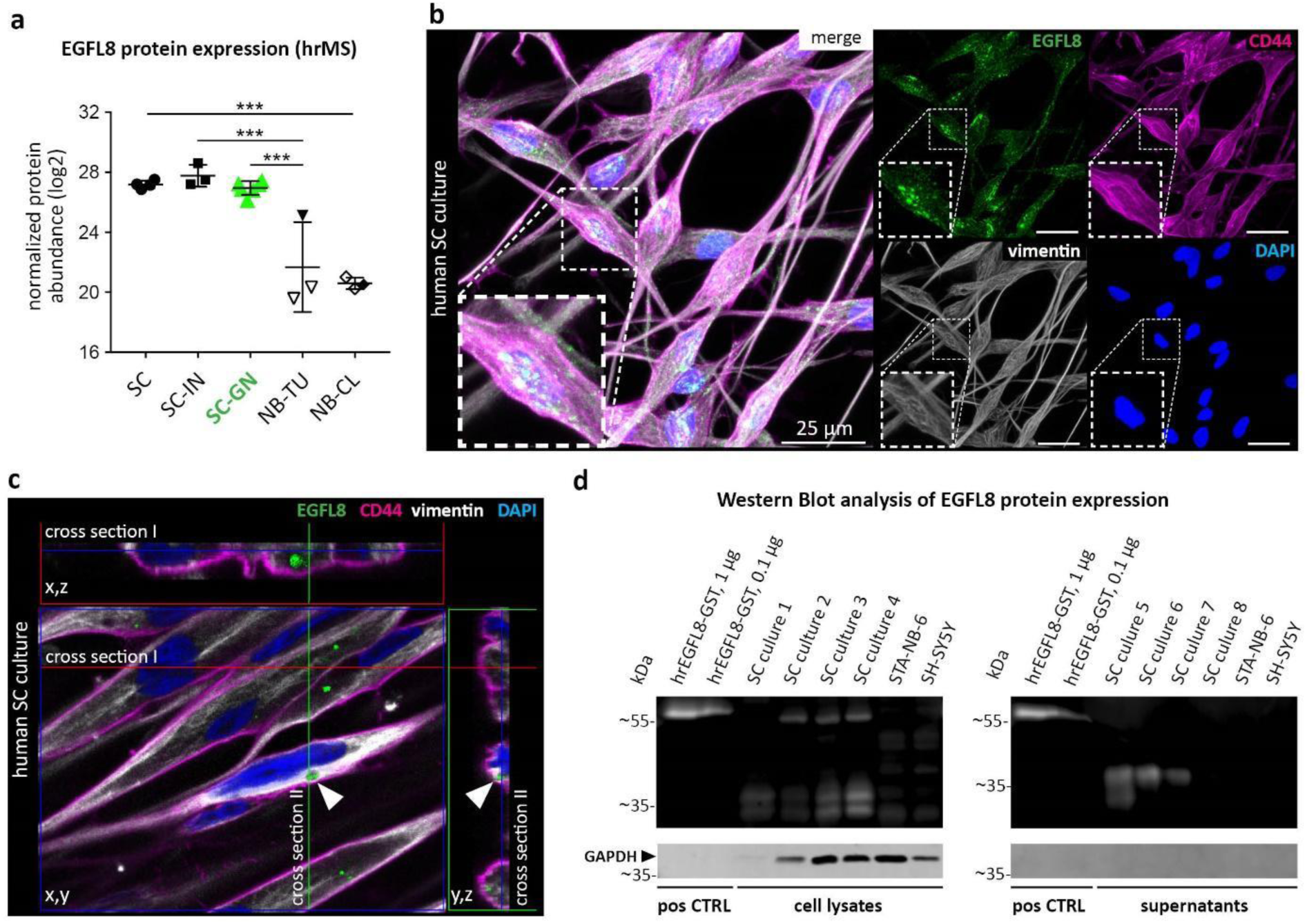
EGFL8 protein expression analysis in tissues containing repair and stromal SCs as well as primary repair SC cultures. **(a)** EGFL8 protein expression levels in primary repair SCs (SC, n=5), injured nerve fascicle tissue (SC-IN, n=3), SC stroma rich ganglioneuromas (SC-GN, n=6), neuroblastomas (NB-TU, n=3) and neuroblastoma cell lines (NB-CL, n=3); lined symbols indicate *MYCN* non-amplified NB-TUs and NB-CLs; *** p ≤ 0.001. **(b)** IF staining of EGFL8, intermediate filament vimentin, surface marker CD44 and DAPI on grown SCs; the enlargement shows a cluster of positive EGFL8 signals. **(c)** 3D analysis of grown SCs confirm EGFL8 positive signals beneath the CD44 positive cell membrane. Arrows and arrowheads indicate individual EGFL8 positive vesicular-like structures in cross sections and clipped image. **(d)** Western blots show EGFL8 protein bands in primary repair SC lysates (n=4) and supernatants (n=4) but not in NB cell line STA-NB-6 and SH-SY5Y lysates and supernatants. Human recombinant (hr) EGFL8 (32kDa) with a GST-tag (26 kDa) was used as positive control. Note that EFGL8 was detected via chemiluminescence, whereas GAPDH was detected via immunofluorescence.

### The EGFL8 protein is secreted by repair Schwann cells *in vitro*

In order to validate EGFL8 expression and to determine its subcellular location in SCs, we stained primary repair SC cultures for EGFL8, CD44 (a cell surface protein strongly expressed by SCs) and VIME. EGFL8 showed an intracellular staining pattern with accumulation of positive signals in clusters of different sizes surrounded by VIME filaments (**Fig. 7b**). 3D analysis illustrated EGFL8 positive vesicular structures embedded within the SC cytoplasm beneath the CD44 positive cell membrane (**Fig. 7c**). In addition, we performed WB analysis for EGFL8 on cell lysates of human primary repair SCs, STA-NB-6 and SH-SY5Y cell lines, and conditioned culture medium (supernatants) of respective cultures. A GST-tagged recombinant EGFL8 protein was used as positive control. EGFL8 has an expected mass of 32 kDa, accordingly, the antibody detected the GST-tagged (GST corresponding to 26 kDa) recombinant EGFL8 protein at around 58 kDa in positive controls (**Fig. 7d**). In all SC samples derived from whole cell lysates, 2 bands were visible at around 32 and 37 kDa. In addition, three SC samples showed an additional band at around 55 kDa. In the SC supernatants, prominent bands were detected at 37 kDa in three out of four SC supernatants, which could indicate that both the intracellular as well as the secreted EGFL8 protein underwent posttranslational modifications.

In line with the predicted secretion of EGFL8, necessary to facilitate its function as neuritogen (see Fig. 6), we show that EGFL8 is stored in vesicular structures within the cytoplasm and secreted by human repair SCs *in vitro*.

## DISCUSSION

This study presents a comparative analysis of human repair SCs in injured nerves and stromal SCs in GNs that builds upon previous efforts to delineate the role of SCs in nerve regeneration and the tumor microenvironment ^1, 13, 27^. By investigating human tissues and primary cultures with deep RNA-sequencing, high-resolution mass spectrometry, confocal imaging, and functional assays, we reveal a similar cellular state and overlapping functional competences of repair SCs and stromal SCs. Our comprehensive approach is the first to identify EGFL8 as a novel SC-secreted neuritogenic factor, highlighting matricellular proteins as tissue active components involved in regenerative and pathological responses of SCs in the peripheral nervous system. Focusing on the interaction of tumor cells and SCs, we developed a co-culture model combined with a flow cytometry-based read-out demonstrating that NB cells react to repair SCs in a similar fashion as peripheral neurons upon injury. Moreover, the established co-culture model is broadly applicable and contributes to the ongoing research in the field of regenerative medicine as well as cancer research aiming to elucidate the interplay of human primary SCs with different cell populations.

### The Schwann cell stroma in ganglioneuroblastomas/ganglioneuromas – an expression of Schwann cell plasticity

The development of mostly benign-behaving GNBs/GNs is hallmarked by an increasing stromal SC population and tumor cell differentiation along the sympathetic neuronal lineage. Since previous studies demonstrated that stromal SCs unlikely descend from tumor cells ^1, 24, 26^, we aimed to understand their origin and cellular state. The accumulation of publications supporting SCs as a highly plastic cell type urged us to investigate whether this reactive/adaptive potential plays a role in GNB/GN development.

The inherent SC plasticity is impressively demonstrated after peripheral nerve injury, where adult SCs undergo substantial expression changes to adapt their cellular functions to the needs of nerve repair, as has been shown by us and others ^13, 36, 43^. In this study, our transcriptomic data reveal that stromal SCs in GNs and repair SCs in damaged nerves share a similar expression profile and nerve repair-associated functions. This finding assigns the cellular state of stromal SCs in GNs to SCs that underwent a phenotypical switch as occurring after nerve damage. The SC stroma development in peripheral neuroblastic tumors indeed exhibits parallels to the nerve injury-induced transformation of adult SCs into a repair cell. Upon nerve injury, adult SCs react to injured axons by cellular reprogramming, which induces the re-expression of genes associated with precursor/immature SC characteristics. This enables SCs to re-enter the cell cycle and to acquire a mesenchymal-like cell state with increased migratory capacity ^2, 44^. These functions match the morphological observation of stromal SCs entering tumors through migration along blood vessels and connective tissue septa and the augmentation of SC stroma over time ^24^. The presence of genes characterizing a pre-myelin developmental stage, e.g. *NGFR, GFAP, ERBB3, CADH19*, SOX2 and *ZEB2*, in SC stroma could also explain why the long axonal processes of ganglionic-like tumor cells in GNs are not myelinated by accompanying stromal SCs ^22^. For example, over-expression of transcription factor Sox2 was shown to block myelination during nerve development and after sciatic nerve crush injury in mice ^45^. Interestingly, the sustained expression levels of Sox2 also correlated with an increased number of macrophages in injured nerves ^45^. Another transcription factor we found highly expressed by stromal SCs was *JUN*, a key modulator that governs the successful transformation into the repair SC identity ^2, 36^. It is important to note that the repair SC state in injured nerves differs from SCs during development. Repair SCs represent a distinct SC phenotype hallmarked by the acquisition of nerve repair-specific functions, such as myelin clearance, macrophage recruitment, upregulation of MHC-II, and the formation of regeneration tracks that support axon re-growth and pathfinding ^9, 10, 12, 13^. Of note, we here identified that genes and pathways associated with these repair functions are highly enriched in stromal SCs. Although we cannot exclude a contribution of mesenchymal stem cells or other precursors able to differentiate into Schwann-like stromal cells during GNB/GN development, the nerve repair-associated expression signature of stromal SCs argue for a repair-like phenotype of stromal SCs, similar to repair SCs emerging in damaged nerves.

The presence of repair-like SCs in GNs also implicates that the tumor cells express factors able to induce and maintain a repair-like SC state in the microenvironment. Thus, we propose that stromal SCs originate from adult SCs that react to peripheral neuroblastic tumor cells in a similar way as to injured neurons. Moreover, the progressing death of ganglionic-like tumor cells and resulting axon degeneration observed in GNBs/GNs ^46^ could supply stromal SCs with cues that trigger the repair-like state and explain why it does not diminish over time. As a consequence, stromal SCs could continuously exert nerve repair-associated functions in the microenvironment that are responsible for a benign tumor development.

### Schwann cell stroma as an effector of neural repair in ganglioneuroblastomas/ganglioneuromas

Recognizing stromal SCs as possible facilitators of nerve repair-associated functions in the tumor microenvironment prompts the question how these functions could affect the behavior of tumor cells. We here show that stromal SCs share the expression of several neurotrophins and axon-guiding proteins with repair SCs in damaged nerves. Hence, neuronal differentiation-inducing cues derived from stromal SCs could be responsible for the differentiation of tumor cells into ganglionic-like cells during GNB/GN development. Proof of concept was provided in functional co-culture experiments, where we exposed five genetically diverse neuroblastoma cell lines, derived from aggressive high-risk NBs, to human primary repair SCs. Both, the direct contact to repair SCs and the in-direct contact to the repair SCs’ secretome, were sufficient to induce neuronal differentiation and to impair proliferation of NB cells. Of note, this anti-tumor effect could be replicated by replacing SCs with recombinant neurotrophic factors discovered within the repair/stromal SC secretome.

In addition to their influence on tumor cells, stromal SCs also hold a considerable potential to modulate the tumor microenvironment. We identified that stromal SCs express MHC-II receptors and chemokines, and confirmed the presence of immune cells in GN sections, which is in line with the increasing reports about the immunomodulatory potential of SCs during nerve regeneration and peripheral neuropathies ^13, 47, 48, 49, 50, 51, 52^. Furthermore, the shared expression signature of stromal/repair SCs contained basement membrane components and ECM remodelers such as metalloproteinases and matricellular proteins. Stromal SCs could therefore recruit and interact with immune cells, as well as execute tissue remodeling functions in the tumor environment with the original goal to rebuild an organized nerve structure similar to repair SCs upon nerve injury.

Taken together, the nerve repair-like phenotype equips stromal SCs with different strategies to influence their environment. Stromal SCs could either directly induce neuronal differentiation of peripheral neuroblastic tumor cells or indirectly manipulate the tumor microenvironment via immunomodulation and ECM remodeling responsible for a favorable tumor development.

### Stromal and repair-type Schwann cells express EGFL8, a matricellular protein of novel neuritogenic function

Our transcriptome and proteome analysis demonstrated a high expression of the matricellular protein EGFL8 in repair SCs and stromal SCs. Moreover, *EGFL8* expression in peripheral neuroblastic tumors correlated with an increased patient survival. We further provided first evidence for a neuritogenic function of human EGFL8, a protein of so far unknown function, as its recombinant form induced neuronal differentiation of NB cell lines at similar efficacy as NGF. This finding underlines the increasingly recognized impact of matricellular proteins in injury response and pathological conditions. Matricellular proteins are dynamically expressed, non-structural ECM proteins that sequester or modulate proteins and growth factors or directly bind to signaling receptors influencing a variety of cellular behaviors ^53^. Their expression is specifically induced during development and upon injury, and plays an important role in tissue remodeling and inflammatory processes ^54, 55, 56^. Stromal SCs and repair SCs also shared the expression of other matricellular proteins such as *SPARC, SPP1* (osteopontin) and *CCN3* (*NOV*). Notably, SC stroma-derived SPARC was previously reported to suppress NB progression by inhibiting angiogenesis and introducing changes in the ECM composition ^57^. We suggest that stromal/repair SCs are a source of various matricellular proteins and actively participate in tissue remodeling events to foster neuronal differentiation. It remains to be evaluated how EGFL8 exerts its neuritogenic effect.

### Exploiting Schwann cell plasticity in therapeutic approaches for aggressive neuroblastomas

The plastic potential of SCs is a double-edged sword. While essential for nerve repair, recent studies point out its adverse effect in neuropathies and epithelial cancer progression ^50, 58^. Here, we demonstrate a favorable impact of SC plasticity on peripheral neuroblastic tumor cells as it manifests in SC stroma during the development of benignly behaving GNB/GN. The cellular similarities between stromal and repair SCs suggest that stromal SCs are able to exert nerve repair-associated functions in the tumor microenvironment. Consequently, the benign tumor development may display the attempt of SCs to ‘repair’ tumor cells and re-establish a nervous structure. Exploiting the strategies repair/stromal SCs use to generate a neuronal (re-)differentiation supporting environment could therefore hold a valuable therapeutic potential.

The prerequisite for a possible treatment approach is the susceptibility of aggressive NBs to SCs. We and others have previously investigated the effect of SCs and their secreted factors on aggressive NB cell lines. These studies confirmed that SCs are able to induce neuronal differentiation and impair the growth of NB cells, which were derived from SC-stroma poor high-risk NBs ^27, 28, 30, 31, 59, 60, 61^. The confirmation that aggressive NB cells, although lacking the ability to attract SCs, are still responsive to SCs, allows elaborating two treatment strategies ^19^. On the one hand, existing therapies could be improved by including SC-derived factors as anti-tumor agents ^1, 62^. On the other hand, understanding how tumor cells induce and maintain a repair SC state in benignly behaving GNB/GN could enable a therapeutic induction of SC stroma in aggressive NBs. Furthermore, identifying how the repair SC state can be sustained is also of high value for the field of regenerative medicine, since one of the main reasons for axonal regeneration failure after injury is the deterioration of repair SCs over time ^63^. Thus, the more detailed knowledge about the molecular processes involved in GNB/GN development and nerve regeneration is promising to enrich treatment approaches for both nerve repair and aggressive NBs.

## CONCLUSION

Our study demonstrates that the cellular state of stromal SCs in GN shares key features with repair SCs in injured nerves. This finding provides essential insights into GNB/GN development as it suggests that the inherent plasticity allows SCs to react to peripheral neuroblastic tumor cells in a similar way as to injured neurons. As a consequence, stromal SCs could exert repair-associated functions that shape an anti-tumor microenvironment and induce the neuronal differentiation of tumor cells responsible for a benign tumor behavior. Using the example of EGFL8, we show that stromal and repair SCs indeed use similar mechanisms to promote neuronal differentiation, which hold considerable treatment possibilities for the therapy of aggressive NBs.

## METHODS

### Human material

The collection and research use of human peripheral nerve tissues and human tumor specimen was conducted according to the guidelines of the Council for International Organizations of Medical Sciences (CIOMS) and World Health Organisation (WHO) and has been approved by the local ethics committees of the Medical University of Vienna (EK2281/2016 and 1216/2018).

### Human peripheral nerve explants and primary Schwann cell cultures

Human peripheral nerves were collected during reconstructive surgery, amputations or organ donations of male and female patients between 16 and 70 years of age. The *ex vivo* nerve injury model as well as the isolation procedure and culture conditions of primary human SCs have been performed as previously described ^13, 64^. Briefly, fascicles were pulled out of nerve explants and digested overnight. The fascicle-derived cell suspension was seeded on PLL/laminin coated dishes and cultured in *SC expansion medium (SCEM:* MEMα GlutaMAX™, 1% Pen/Strep, 1 mM sodium pyruvate, 25 mM HEPES, 10 ng/mL hu FGF basic, 10 ng/mL hu Heregulin-β1, 5 ng/mL hu PDGF-AA, 0.5% N2 supplement, 2 µM forskolin and 1% FCS. Cells of the initial seeding represent passage 0 (p0). Half of the medium was changed twice a week. When the cultures reached approx. 80% confluence, contaminating FBs were depleted by exploiting their ability to adhere more rapidly to plastic. Enriched passage 1 (p1) SC cultures of about 96% purity, as determined via positivity for the SC marker S100B, were used for experimentation. For the *ex vivo* nerve injury model, about 1.5 cm long human nerve fascicles were subjected to an *ex vivo* degeneration period of 8 days in SCEM + 10% FCS at 37°C (= injured nerve fascicle). During that time, axons degenerate and SCs adapt the repair phenotype within the explant ^13^.

### Neuroblastoma/ganglioneuroma tissue and neuroblastoma cell lines

Tumor specimen from diagnostic NB tumors (NB-TU, n=18) and GN tumors (GN-TU, n=6) have been collected during surgery or biopsy for diagnostic purposes and left-overs were cryopreserved until analysis. Cryosections of GN tissue were analyzed for SC stroma rich areas identified by H+E-staining, immmunofluorescence staining for SC marker S100B, and confirmed by a pathologist. The corresponding tumor region was excised using a scalpel blade and cryopreserved until RNA and protein extraction.

The used NB cell lines are derived from biopsies or surgical resection of aggressively behaving NB tumors of patients suffering from high-risk metastatic NBs. In-house established, low passage NB cell lines STA-NB-6, −7 −10 and −15 as well as the NB cell lines SK-N-SH, SH-SY5Y, IMR5 and CLB-Ma were cultured in *MEMα complete* (MEMα GlutaMAX™, 1% Pen/Strep, 1 mM sodium pyruvate, 25 mM HEPES and 10% FCS). The NB cell lines and NBs differ in their genomic background including *MYCN*-amplification status. An overview of NB cell line as well as NB and GN tumor characteristics is provided in **Supplementary (S.) Tables 1, 2 & 3**, respectively.

### The co-culture model of primary Schwann cells and neuroblastoma cell lines

Enriched human p1 SCs from at least 3 independent donors were co-cultured with 5 NB cell lines (STA-NB-6, STA-NB-10, IMR5, SH-SY5Y and CLB-Ma), respectively. First, SCs were seeded in PLL/laminin coated wells of a 6-well plate in SCEM. At day 1 and day 2, half of the media was exchanged with MEMα complete. At day 3, total media was changed to MEMα complete and NB cells were seeded directly to the p1 SC cultures as well as in PLL/laminin coated trans-wells (24 mm Inserts, 0.4 µm polyester membrane, COSTAR) placed above SC cultures, alongside with respective controls. Two third of the media was changed twice a week and one day prior to FACS analyses on day 8 and day 16.

For IF analysis, SCs from 3 independent donors were co-cultured with STA-NB-6 or CLB-Ma NB cell lines in coated wells of an 8-well chamber slide (Ibidi), respectively, alongside with controls for 11 days.

### Proliferation and differentiation FACS panels

All antibody details are listed in **S.Table 3**. If not stated otherwise, all steps of the staining procedures were performed on ice. The following antibodies have been conjugated to fluorochromes using commercially available kits according to the manufacturer’s instructions: anti-S100B has been conjugated to FITC (FLUKA) using Illustra NAP-5 columns (GE Healthcare), anti-GD2 (ch14:18, kindly provided by Professor Rupert Handgretinger, Department of Hematology/Oncology, Children’s University Hospital, Tübingen, Germany) has been conjugated to AF546 using the AlexaFluor® 546 protein labeling kit (Molecular probes) and anti-NF200 has been conjugated to AF647 using the AlexaFluor® 647 protein labeling kit (Molecular probes).

Cells were detached using Accutase (LifeTechnologies) and washed with FACS-buffer (1x PBS containing 0.1% BSA and 0.05% NaAzide). For the *differentiation FACS panel*, cells were incubated with GD2-AF546 for 20 min, washed once with FACS-buffer and fixed using Cytofix/Cytoperm (BD Biosciences) in the dark for 20 min. After washing with 1x perm/wash (BD), cells were stained with anti-S100B-FITC and NF200-A647 for 20 min. Cells were washed in 1x perm/wash and analyzed immediately at the FACSFortessa flow cytometer equipped with the FACSDiva software (both BD). For the *proliferation FACS panel*, 1 µM EdU was added to cultures for about 15 hours. Cells were detached, washed and fixed in Roti-Histofix 4% for 20 min at RT. Permeabilization and EdU detection was carried out using the Click-iT EdU Alexa Fluor 647 Flow Cytometry Assay Kit (Thermo Fisher Scientific) according to the manufacturer’s manual. Additional extracellular/intracellular staining was performed with GD2-A546 and anti-S100B-FITC antibodies in 1x saponin-based perm/wash for 30 min. After washing, cells were resuspended in 1x saponin-based perm/wash, 1 μl of FxCycle Violet (LifeTechnologies) DNA dye was added and samples were analyzed immediately at the FACSFortessa.

### Immunofluorescence staining and confocal image analysis

All antibody details are listed in **S.Table 3**. If not stated otherwise, the staining procedure was performed on RT and each washing step involved three washes with 1x PBS for 5 min. Primary antibodies against extracellular targets were diluted in 1x PBS containing 1% BSA and 1% serum; primary antibodies against intracellular targets were diluted in 1x PBS containing 1% BSA, 0.1% TritonX-100 and 1% serum. Briefly, co-cultures were fixed with Roti-Histofix 4% (ROTH) for 20 min at 4°C, washed, and blocked with 1x PBS containing 1% BSA and 3% serum for 30 min. Cells were incubated with primary antibodies against extracellular targets, washed and incubated with appropriate secondary antibodies for 1h. Samples were then again fixed with Roti-Histofix 4% for 10 min. After washing, cells were permeabilized and blocked with 1x PBS containing 0.3% TritonX-100 and 3% serum for 10 min. When required, TUNEL staining was performed after permeabilization according to the manufacturer’s protocol (PROMEGA). Samples were then incubated with primary antibodies against intracellular targets, washed and incubated with the appropriate secondary antibodies for 1h. Finally, cells were incubated with 2 µg/mL DAPI in 1x PBS for 2 min, washed and embedded in Fluoromount-G mounting medium (SouthernBiotech). Images were acquired with a confocal laser scanning microscope (LEICA, TCS SP8X) using Leica LAS AF software. Confocal images are shown as maximum projection of total z-stacks and brightness and contrast were adjusted in a homogenous manner using the Leica LAS AF software.

### RNA isolation, RNA sequencing and gene expression analysis

Fresh frozen SC stroma-rich areas derived from diagnostic GNs (SC-GN, n=6) were homogenized with the gentleMACS Dissociator (Miltenyi) using 1 mL of TRIzol per sample and the predefined RNA-01 gentleMACS program. RNA isolation was performed with the miRNeasy micro kit (Qiagen) following the manufacturer’s protocol. Quantity and integrity of extracted RNA were assessed by the Qubit RNA HS Assay Kit (Life Technologies) and the Experion RNA StdSens Assay Kit (BioRad), respectively. 30 ng total RNA (RQI≥8) was used for library preparation following the NEBNext Ultra RNA Library Prep Kit for Illumina protocol (New England BioLabs) with the Poly(A) mRNA Magnetic Isolation Module (New England BioLabs). After cDNA synthesis, the library was completed in an automated way at the EMBL Genomics Core Facility (Heidelberg, Germany). RNA-Seq was performed at the Illumina HiSeq 2000 platform and 50 bp-single-end reads were generated.

The generated data were bioinformatically analyzed together with our previously published transcriptomic data sets of human primary SCs (SC, n=5), human injured fascicle explants (SC-IN, n=3) and NB cell lines STA-NB-6 (in triplicates), STA-NB-7 and STA-NB-15 (NB-CL, n=3) ^13^, and diagnostic, untreated stage 4 NBs (NB-TU, n=15) ^65^. Respective GEO identifiers can be found in **S.Table 4**.

Short read sequencing data was quality checked using FASTQC (http://www.bioinformatics.babraham.ac.uk/projects/fastqc) and QoRTs ^66^ and then aligned to the human genome hs37d5 (ftp://ftp.1000genomes.ebi.ac.uk/) using the STAR aligner ^67^ yielding a minimum of 11.6 million aligned reads in each sample. Further analysis was performed in R statistical environment using Bioconductor packages^68^. Count statistics for Ensembl (GRCh37.75) genes were obtained by the “featureCounts” function (package “Rsubread”) and differential expression analysis was performed by edgeR and voom ^69, 70^. For differential gene expression analysis only genes passing a cpm (counts per gene per million reads in library) cut-off of 1 in more than two samples were included. All p-values were corrected for multiple testing by the Benjamini-Hochberg method. Genes with an adjusted q-value <0.05 and a log2 fold change > 1 (|log2FC|>1) were referred to as ‘significantly regulated’ and used for functional annotation analysis via DAVID database ^71^.

### Protein isolation, high-resolution mass spectrometry and expression analysis

Fresh frozen diagnostic GN-derived SC stroma-rich areas (SC-GN, n=6), diagnostic high-risk NB tumors (NB-TU, n=3) as well as low-passage NB cell lines STA-NB-7, STA-NB-2 and STA-NB-10 (NB-CL, n=3) were used for proteomic analysis (see **S.Table 1 & 2** for tumor and cell line characteristics). Protein isolation from cells and tissue, mass spectrometry sample preparation and liquid chromatography-mass spectrometry (LC-MS) has been carried out as described previously ^13, 72^, all samples were measured in two technical replicates.

Label-free quantitative data analysis was performed using MaxQuant 1.3.0.5 with the Andromeda search engine and the Perseus statistical analysis package ^73, 74^. The data were analysed together with previously generated proteomic data set of human primary SCs (SC, n=4) and human injured fascicle explants (SC-IN, n=3) ^13^. The mass spectrometry proteomics data have been deposited to the ProteomeXchange Consortium (http://proteomecentral.proteomexchange.org) via the PRIDE partner repository ^75^ with the dataset identifier PXD018267.

### Western Blot analysis

All antibody details are listed in **S.Table 3**. Western blot analysis was performed as previously described ^76, 77^. 1x TBS-T was used for all washing steps that were performed three times for 5 min after each antibody incubation. Briefly, frozen cell aliquots were thawed, pelleted and lysed by addition of RIPA buffer. Culture media were centrifuged at 300 g for 10 min at 4°C to remove cellular debris. The supernatants were mixed with −20°C EtOH (1:5), precipitated at −20°C for 20 h, centrifuged at 4000xg for 40 min at 4°C and the dried pellet was lysed by addition of RIPA buffer. Protein extracts were stored in Protein LoBind tubes (Eppendorf) at −80°C. Protein concentrations were determined via Bradford assay (BioRad). Protein extracts were mixed with SDS-loading buffer, denatured for 5 min at 95°C, separated on a 10% SDS/PAA gel and blotted onto methanol-activated Amersham Hybond-P PVDF membranes. Membranes were blocked using 1x T-BST with 5% w/v nonfat dry milk for 30 min and incubated with anti-EGFL8 followed by HRP-conjugated secondary antibody. The blots were developed using the WesternBright Quantum detection kit (Advansta) and visualized with the FluorChemQ imaging system (Alpha Innotech, San Leandro, USA). Subsequently, membranes were incubated with anti-GAPDH followed by IRdye680T labeled secondary antibody. Blots were analyzed using the Odyssey imaging system (Licor) and the Odyssey software v3.0.

### Statistical analyses

If not mentioned otherwise, GraphPad Prism5 was used for statistical analysis. Values were given as means ± SD (n ≥ 3). For parametric analysis, ANOVA and Bonferroni post-hoc test was performed. p-values ≤0.05 were considered significant.

### Data Availability Section

All data sets produced and used in this study are available in public repositories as listed in **S.Table 4**. The mass spectrometry proteomics data have been deposited to the ProteomeXchange Consortium (http://proteomecentral.proteomexchange.org) via the PRIDE partner repository ^75^ with the dataset identifier PXD018267.

## Author contributions

T.W. and S.T.-M. planned experiments, performed research, analyzed and interpreted data and wrote manuscript; H.D., A.B. and F.R. performed research and analyzed data; C.F. and M.K. developed bioinformatics tools and analyzed data; C.G. analyzed and interpreted data; R.W. provided essential material; P.F.A. and I.M.A. conceptualized the project, interpreted data and reviewed manuscript.

## Acknowledgements

This study was supported by Österreichische Forschungsförderungsgesellschaft (FFG) grants (ID:844198, TisQuant, EraSME, by the Austrian Research Promotion Agency, to P.F. Ambros and VISIOMICS, Coin Netzwerke, to S. Taschner-Mandl), the European Union’s Seventh Framework Program (FP7/2007–2013) under the project ENCCA, grant agreement HEALTH-F2-2011-261474, the Herzfeldersche Familienstiftung, Modicell (MC-IAPP Project 285875) and St. Anna Kinderkrebsforschung.

## ABBREVIATIONS

SC: Schwann cell
NT: peripheral neuroblastic tumor
NB: neuroblastoma
GNB: ganglioneuroblatoma
GN: ganglioneuroma
RT: room temperature
IF: immunofluorescence

## Conflict of interest

The authors declare no conflict of interest.

## For More Information

www.science.ccri.at

## SUPPLEMENTARY TABLES

**Supplementary table 1.**
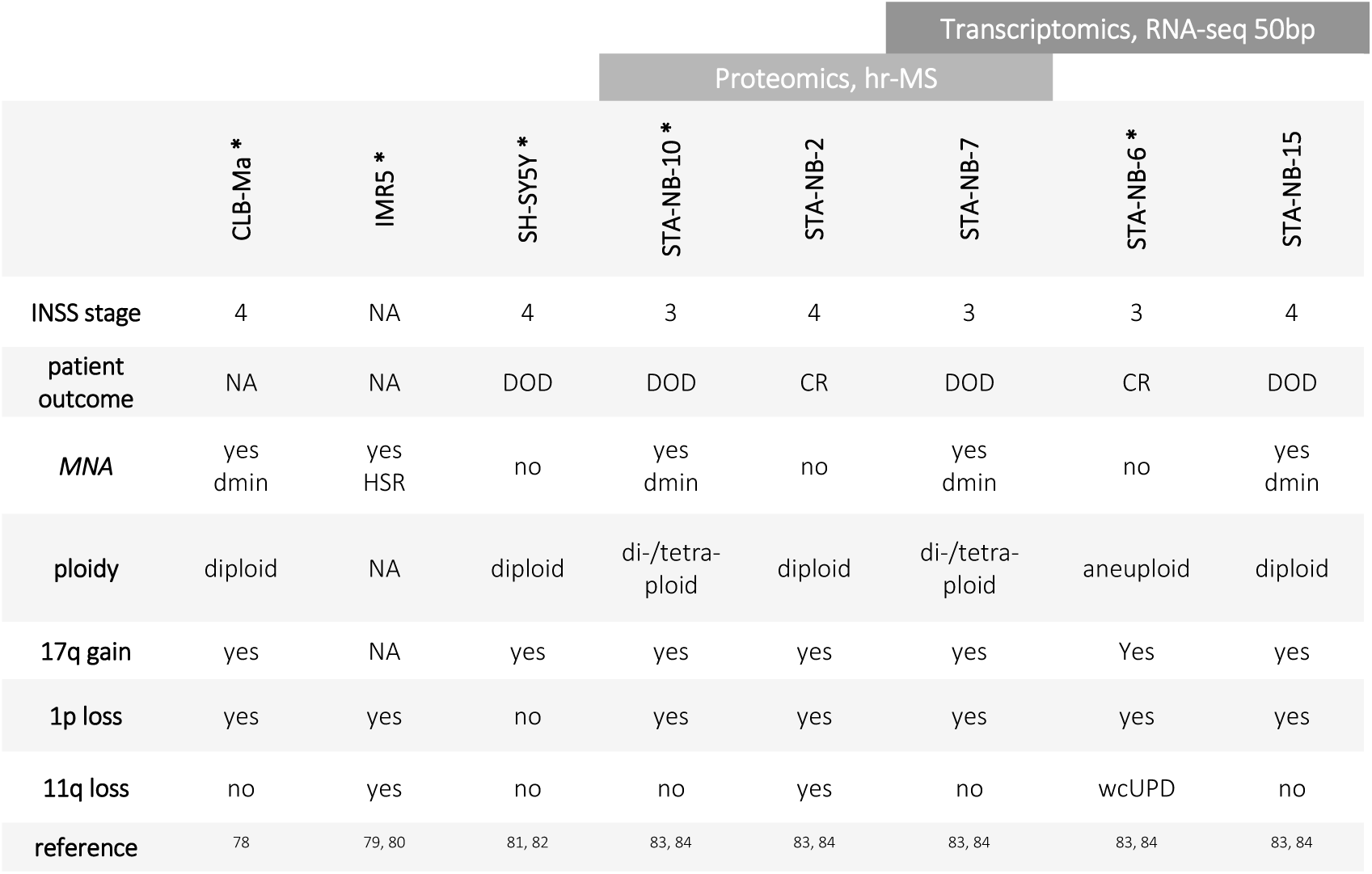
NBT cell lines characteristics. INSS (International Neuroblastoma Staging System), NA (not available), MNA (*MYCN* amplification), HSR (homogeneously staining regions), WT (wild type), DOD (death of disease), CR (complete remission), dmin (double minutes), wcUPD (whole chromosome uniparental disomy), with ‘*’ marked cell lines were used for co-coculture with primary repair SCs

**Supplementary table 2.**
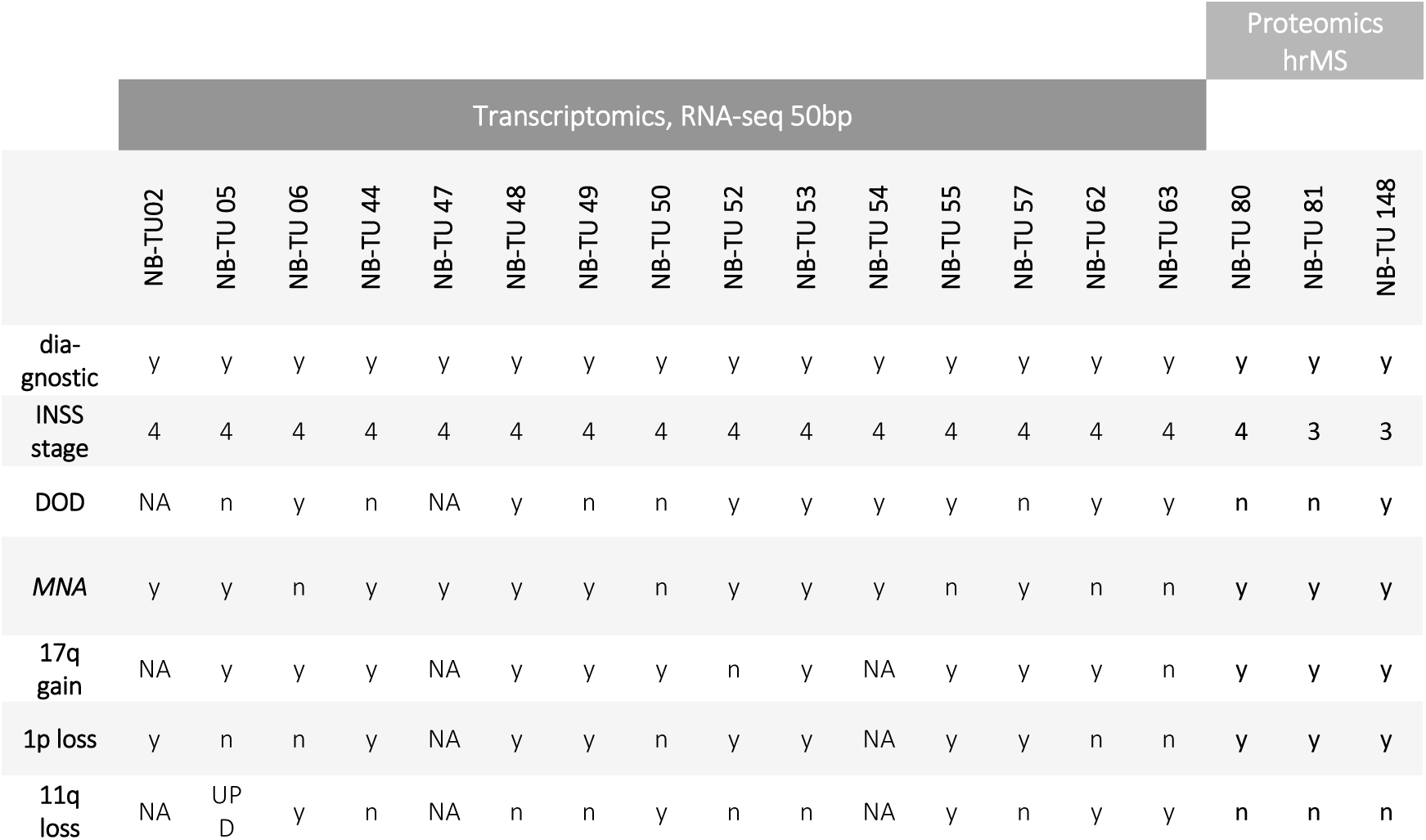
NBT characteristics. INSS (International Neuroblastoma Staging System), y (yes), n (no), NA (not available), MNA (*MYCN* amplification), DOD (death of disease), UPD (uniparental disomy), GEO (Gene Expression Omnibus)

**Supplementary table 3.** Antibody list. *manually labeled, perm = permeabilization necessary, RT = room temperature

| 1<sup>st</sup> antibodies |  |  |  | Application |  |  |  |  |  |
| --- | --- | --- | --- | --- | --- | --- | --- | --- | --- |
| Antigen | Species | catalog No | company | immunofluorescence |  | flow cytometry |  | Western Blot |  |
|  |  |  |  | dilution | comment | dilution | comment | dilution | comment |
| S100B | rabbit | #Z0311 | DAKO | 1:200 | 1 hr, RT, perm |  |  |  |  |
| Ki67 | mouse | NCL-Ki67-MM1 | Leica Microsystems | 1:50 | 1 hr, RT, perm |  |  |  |  |
| CD44 | mouse | #M7082 | DAKO | 1:50 | 1 hr, RT, perm |  |  |  |  |
| vimentin | mouse | #M0725 | DAKO |  |  | 1:200 | 20 min, 4°C, perm |  |  |
| vimentin | chicken | #AB5733 | Merck Millipore | 1:200 | 1 hr, RT, perm |  |  |  |  |
| NF200 | mouse | MAB5266 | Merck Milipore | 1:200 | 1 hr, RT, perm |  |  |  |  |
| NGFR-A647* | rabbit | #8238S | CellSignaling |  |  | 1:60 | 20 min, 4°C perm |  |  |
| S100B-FITC* | rabbit | #Z0311 | DAKO |  |  | 1:50 | 20 min, 4°C, perm |  |  |
| GD2-FITC* | humanized chinese hamster | ch14:18 | POLYMUN GmbH | 1:70 | o.n., 4°C |  |  |  |  |
| GD2-A546* | humanized chinese hamster | ch14:18 | POLYMUN GmbH | 1:70 | o.n., 4°C | 1:1000 | 20 min, 4°C |  |  |
| NF200-A647* | mouse | MAB5266 | Merck Milipore |  |  | 1:400 | 20 min, 4°C, perm |  |  |
| EGFL8 | rabbit | #PA5-63929 | Thermo Scientific | 1:100 | o.n., 4°C, perm |  |  | 1:333 | o.n., 4°C |
| CD3 | mouse | #C7048 | Sigma-Aldrich | 1:50 | o.n., 4°C |  |  |  |  |
| HLA-DR-α1 | mouse | #M0746 | DAKO | 1:50 | o.n., 4°C |  |  |  |  |
| GAPDH | mouse | #32233 | Santa Cruz |  |  |  |  | 1:2000 | 1h, RT |

**Supplementary table 3 | Antibody list**
| 2<sup>nd</sup> antibodies |  |  |  | Application |  |  |  |  |  |
| --- | --- | --- | --- | --- | --- | --- | --- | --- | --- |
| Antigen | Species | catalog No | company | immunofluorescence |  | flow cytometry |  | Western Blot |  |
|  |  |  |  | dilution | comment | dilution | comment | dilution | comment |
| α rb FITC | swine | #F0205 | DAKO | 1:50 | 1 hr, RT |  |  |  |  |
| α ms AF594 | goat | #A11032 | LifeTechnologies | 1:300 | 1 hr, RT | 1:1000 | 20 min, 4°C |  |  |
| α ch AF647 | goat | #SA5-10073 | LifeTechnologies | 1:300 | 1 hr, RT |  |  |  |  |
| α ms IRdye680 | goat | P/N 925-68020 | LI-COR |  |  |  |  | 1:10000 | 1h, RT |
| α rb HRP | goat | #7074 | Cell Signaling |  |  |  |  | 1:1000 | 10 min, RT |

**Supplementary table 4.**
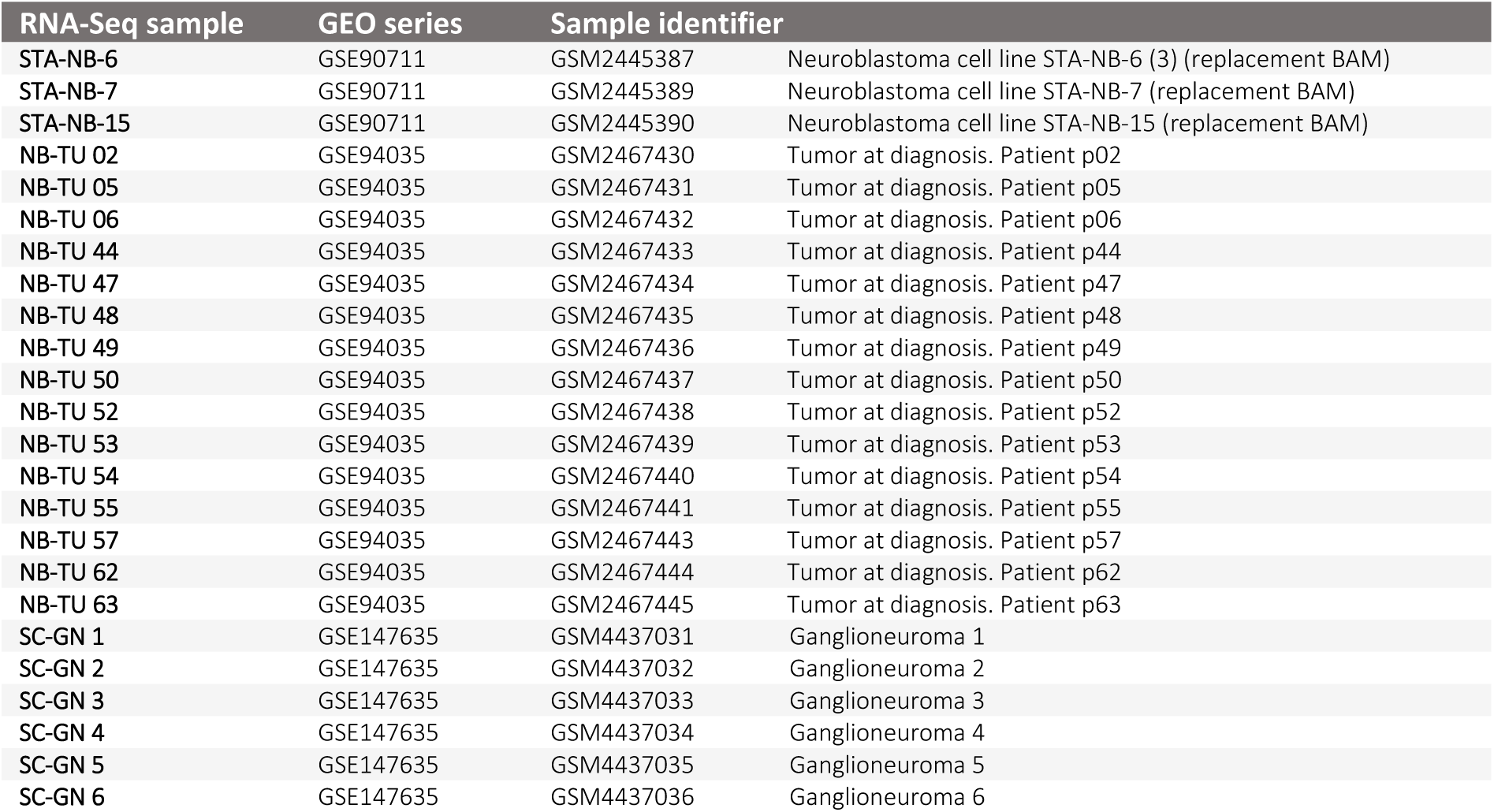
Data repository identifier for samples used in this study. RNA-sequencing datasets were uploaded to the gene expression omnibus (GEO) repository (https://www.ncbi.nlm.nih.gov/geo/)

| RNA-Seq sample | GEO series | Sample identifier |  |
| --- | --- | --- | --- |
| STA-NB-6 | GSE90711 | GSM2445387 | Neuroblastoma cell line STA-NB-6 (3) (replacement BAM) |
| STA-NB-7 | GSE90711 | GSM2445389 | Neuroblastoma cell line STA-NB-7 (replacement BAM) |
| STA-NB-15 | GSE90711 | GSM2445390 | Neuroblastoma cell line STA-NB-15 (replacement BAM) |
| NB-TU 02 | GSE94035 | GSM2467430 | Tumor at diagnosis. Patient p02 |
| NB-TU 05 | GSE94035 | GSM2467431 | Tumor at diagnosis. Patient p05 |
| NB-TU 06 | GSE94035 | GSM2467432 | Tumor at diagnosis. Patient p06 |
| NB-TU 44 | GSE94035 | GSM2467433 | Tumor at diagnosis. Patient p44 |
| NB-TU 47 | GSE94035 | GSM2467434 | Tumor at diagnosis. Patient p47 |
| NB-TU 48 | GSE94035 | GSM2467435 | Tumor at diagnosis. Patient p48 |
| NB-TU 49 | GSE94035 | GSM2467436 | Tumor at diagnosis. Patient p49 |
| NB-TU 50 | GSE94035 | GSM2467437 | Tumor at diagnosis. Patient p50 |
| NB-TU 52 | GSE94035 | GSM2467438 | Tumor at diagnosis. Patient p52 |
| NB-TU 53 | GSE94035 | GSM2467439 | Tumor at diagnosis. Patient p53 |
| NB-TU 54 | GSE94035 | GSM2467440 | Tumor at diagnosis. Patient p54 |
| NB-TU 55 | GSE94035 | GSM2467441 | Tumor at diagnosis. Patient p55 |
| NB-TU 57 | GSE94035 | GSM2467443 | Tumor at diagnosis. Patient p57 |
| NB-TU 62 | GSE94035 | GSM2467444 | Tumor at diagnosis. Patient p62 |
| NB-TU 63 | GSE94035 | GSM2467445 | Tumor at diagnosis. Patient p63 |
| SC-GN 1 | GSE147635 | GSM4437031 | Ganglioneuroma 1 |
| SC-GN 2 | GSE147635 | GSM4437032 | Ganglioneuroma 2 |
| SC-GN 3 | GSE147635 | GSM4437033 | Ganglioneuroma 3 |
| SC-GN 4 | GSE147635 | GSM4437034 | Ganglioneuroma 4 |
| SC-GN 5 | GSE147635 | GSM4437035 | Ganglioneuroma 5 |
| SC-GN 6 | GSE147635 | GSM4437036 | Ganglioneuroma 6 |

**Supplementary table 5.**
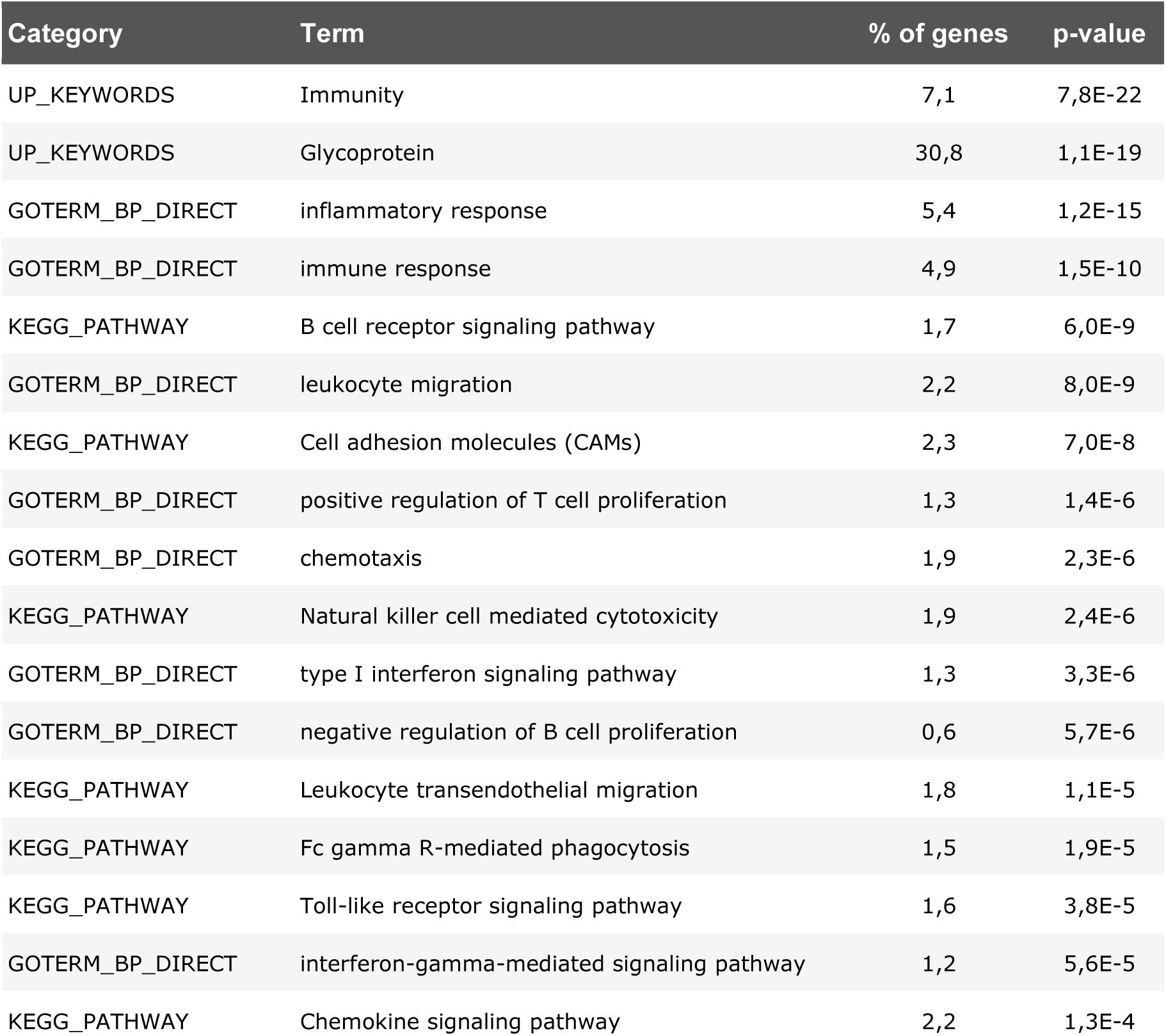
Gene ontology and functional annotation analysis of SC stroma genes not shared with repair SCs. Gene ontology term (GOTERM), cellular compartment (CC), biological process (BP).

| Category | Term | % of genes | p-value |
| --- | --- | --- | --- |
| UP_KEYWORDS | Immunity | 7,1 | 7,8E-22 |
| UP_KEYWORDS | Glycoprotein | 30,8 | 1,1E-19 |
| GOTERM_BP_DIRECT | inflammatory response | 5,4 | 1,2E-15 |
| GOTERM_BP_DIRECT | immune response | 4,9 | 1,5E-10 |
| KEGG_PATHWAY | B cell receptor signaling pathway | 1,7 | 6,0E-9 |
| GOTERM_BP_DIRECT | leukocyte migration | 2,2 | 8,0E-9 |
| KEGG_PATHWAY | Cell adhesion molecules (CAMs) | 2,3 | 7,0E-8 |
| GOTERM_BP_DIRECT | positive regulation of T cell proliferation | 1,3 | 1,4E-6 |
| GOTERM_BP_DIRECT | chemotaxis | 1,9 | 2,3E-6 |
| KEGG_PATHWAY | Natural killer cell mediated cytotoxicity | 1,9 | 2,4E-6 |
| GOTERM_BP_DIRECT | type I interferon signaling pathway | 1,3 | 3,3E-6 |
| GOTERM_BP_DIRECT | negative regulation of B cell proliferation | 0,6 | 5,7E-6 |
| KEGG_PATHWAY | Leukocyte transendothelial migration | 1,8 | 1,1E-5 |
| KEGG_PATHWAY | Fc gamma R-mediated phagocytosis | 1,5 | 1,9E-5 |
| KEGG_PATHWAY | Toll-like receptor signaling pathway | 1,6 | 3,8E-5 |
| GOTERM_BP_DIRECT | interferon-gamma-mediated signaling pathway | 1,2 | 5,6E-5 |
| KEGG_PATHWAY | Chemokine signaling pathway | 2,2 | 1,3E-4 |

**Supplementary table 6.** Gene ontology and functional annotation analysis of injured nerve associated repair SC genes not shared with stromal SCs. Gene ontology term (GOTERM), cellular compartment (CC), biological process (BP)

| Category | Term | % of genes | p-value |
| --- | --- | --- | --- |
| UP_KEYWORDS | Endoplasmic reticulum | 13,3 | 2,4E-26 |
| UP_KEYWORDS | Transport | 18,9 | 6,1E-23 |
| GOTERM_CC_DIRECT | endoplasmic reticulum membrane | 11,1 | 2,0E-20 |
| KEGG_PATHWAY | Protein processing in endoplasmic reticulum | 4,4 | 2,1E-17 |
| UP_KEYWORDS | Protein transport | 7,8 | 3,8E-16 |
| UP_KEYWORDS | Acetylation | 24,7 | 1,2E-13 |
| UP_KEYWORDS | Golgi apparatus | 8,8 | 1,2E-13 |
| GOTERM_BP_DIRECT | response to endoplasmic reticulum stress | 2,0 | 6,9E-9 |
| GOTERM_BP_DIRECT | protein folding in endoplasmic reticulum | 0,6 | 3,6E-4 |
| GOTERM_BP_DIRECT | COPII vesicle coating | 1,8 | 9,0E-9 |
| UP_KEYWORDS | Lysosome | 3,5 | 3,6E-8 |
| GOTERM_BP_DIRECT | oxidation-reduction process | 6,1 | 3,4E-7 |
| GOTERM_BP_DIRECT | protein N-linked glycosylation | 0,8 | 3,8E-3 |

## SUPPLEMENTARY FIGURES AND FIGURE LEGENDS

**Supplementary figure 1.**
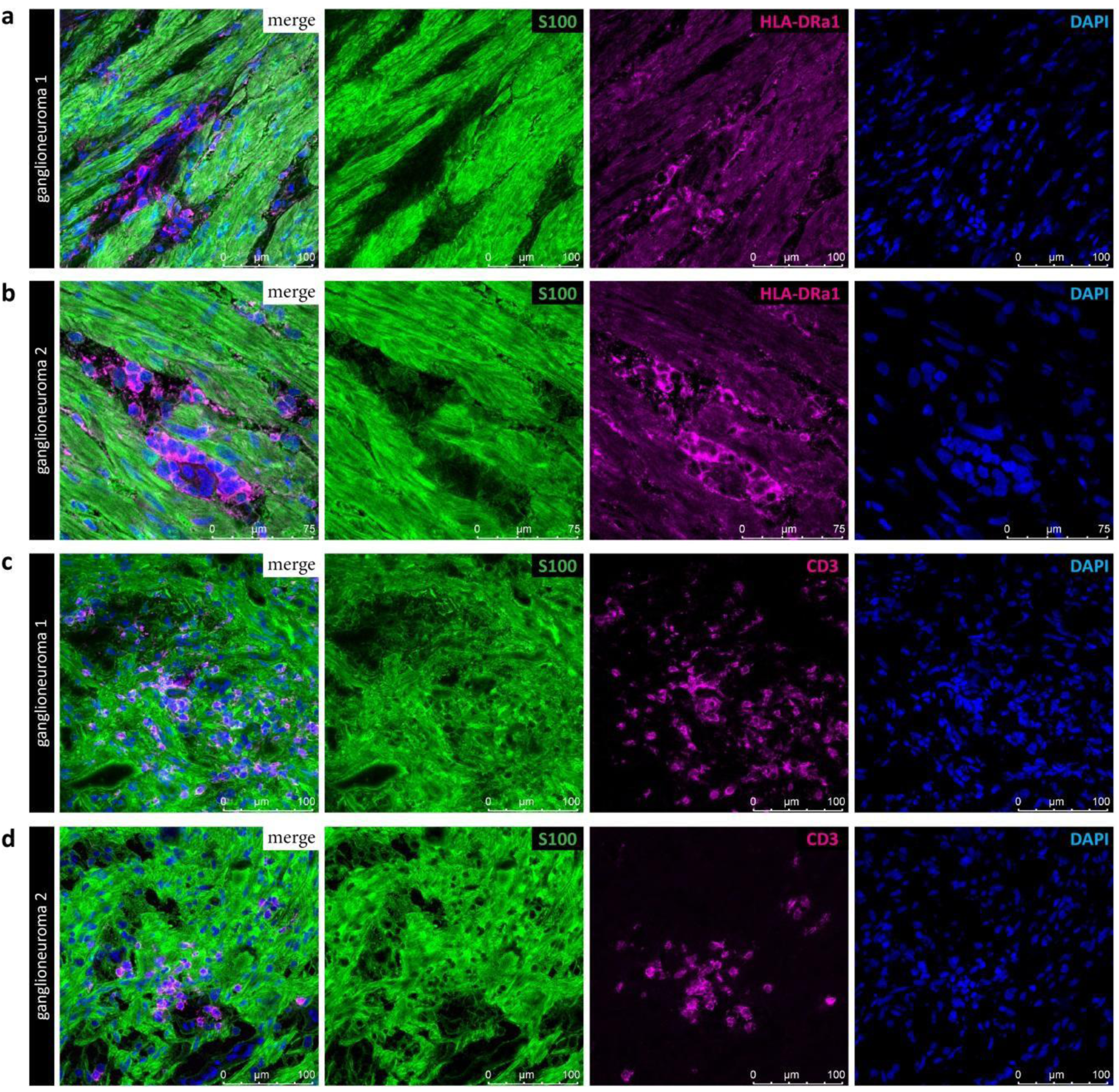
Validation of HLA-DR and CD3 expression in ganglioneuroma. Representative immunofluorescence images of fresh frozen ganglioneuroma sections stained for S100B and HLA-DR **(a-b)** and S100B and CD3 **(c-d)**.

**Supplementary figure 2.**
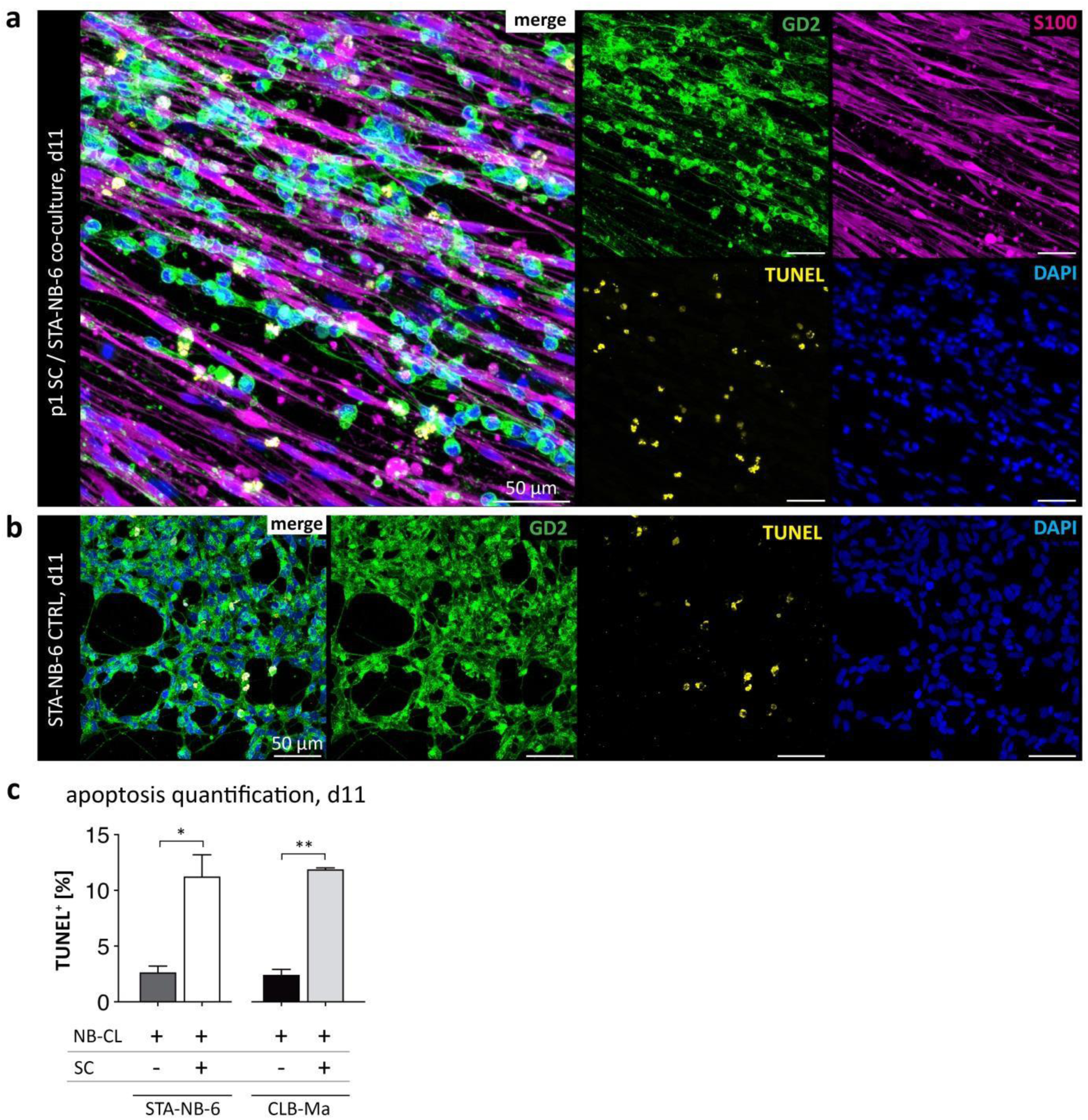
Quantification of apoptotic NBT cells after direct contact to repair SCs *in vitro*. NBT cell lines STA-NB-6 and CLB-Ma were co-cultured with primary SCs for 11 days. IF staining for GD2, S100B and DAPI was combined with a TUNEL assay that detects fragmented DNA typical for apoptotic cells. Representative confocal images of STA-NB-6 **(a)** control cultures and **(b)** co-cultures. **(c)** Bar diagrams show the percentage of GD2^+^/TUNEL^+^ STA-NB-6 and CLB-Ma cells in control (CTRL) and co-cultures (+SCs) at day 11 ± SEM (n≥3); * p ≤ 0.05; ** p ≤ 0.01.

**Supplementary figure 3.**
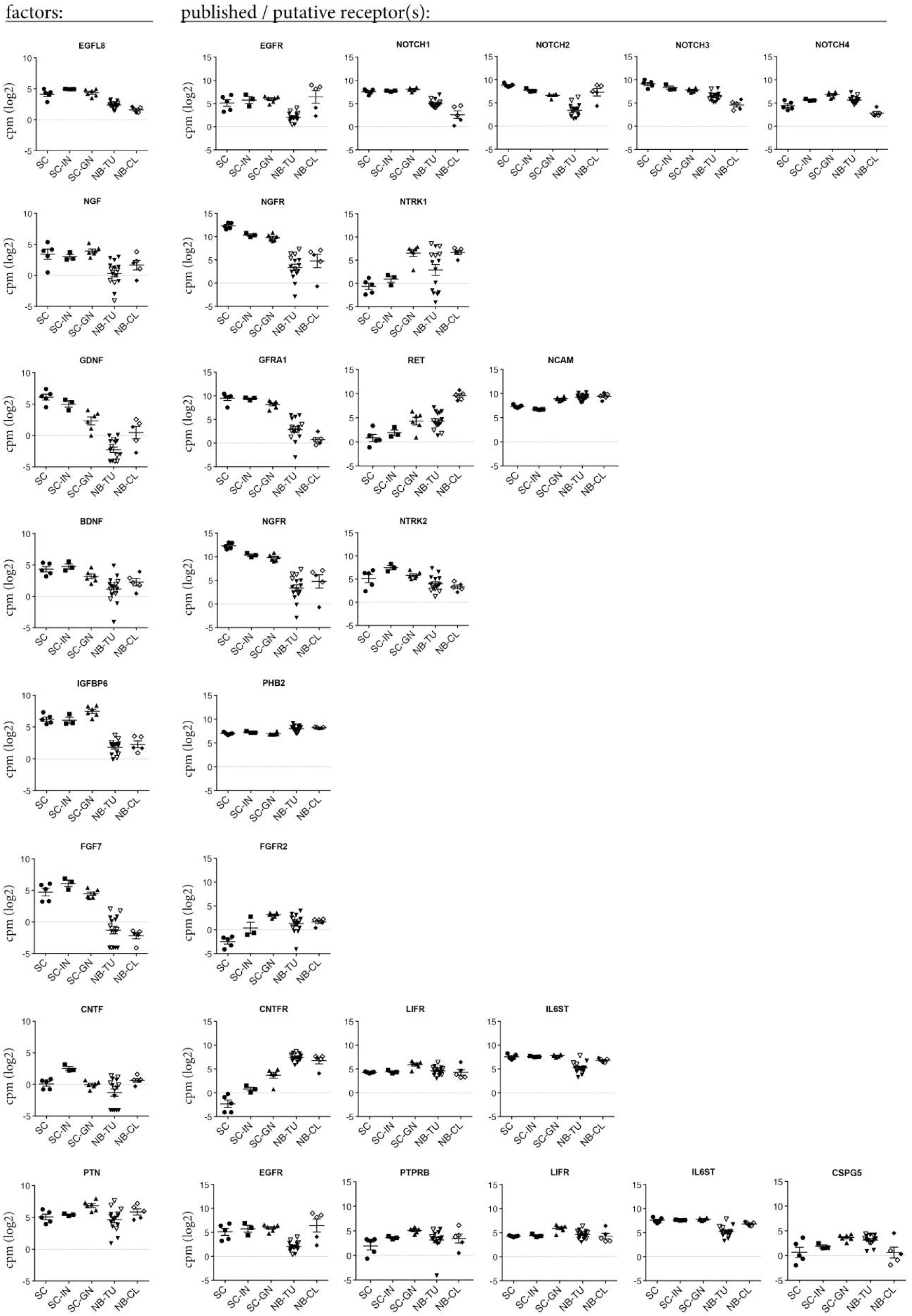
Secreted candidate factors and putative receptors. Expression levels of chosen candidate factors and published or putative receptors in primary repair SCs (SC), injured nerve fascicle tissue (SC-IN), SC stroma rich ganglioneuroma tissue (SC-GN), NBT tissue (NB-TU) and NBT cell lines (NB-CL); lined symbols indicate *MYCN* non-amplified non-amplified NB-TUs and NB-CLs. ** p ≤ 0.01***, p ≤ 0.001, n.s. not significant.

**Supplementary figure 4.**
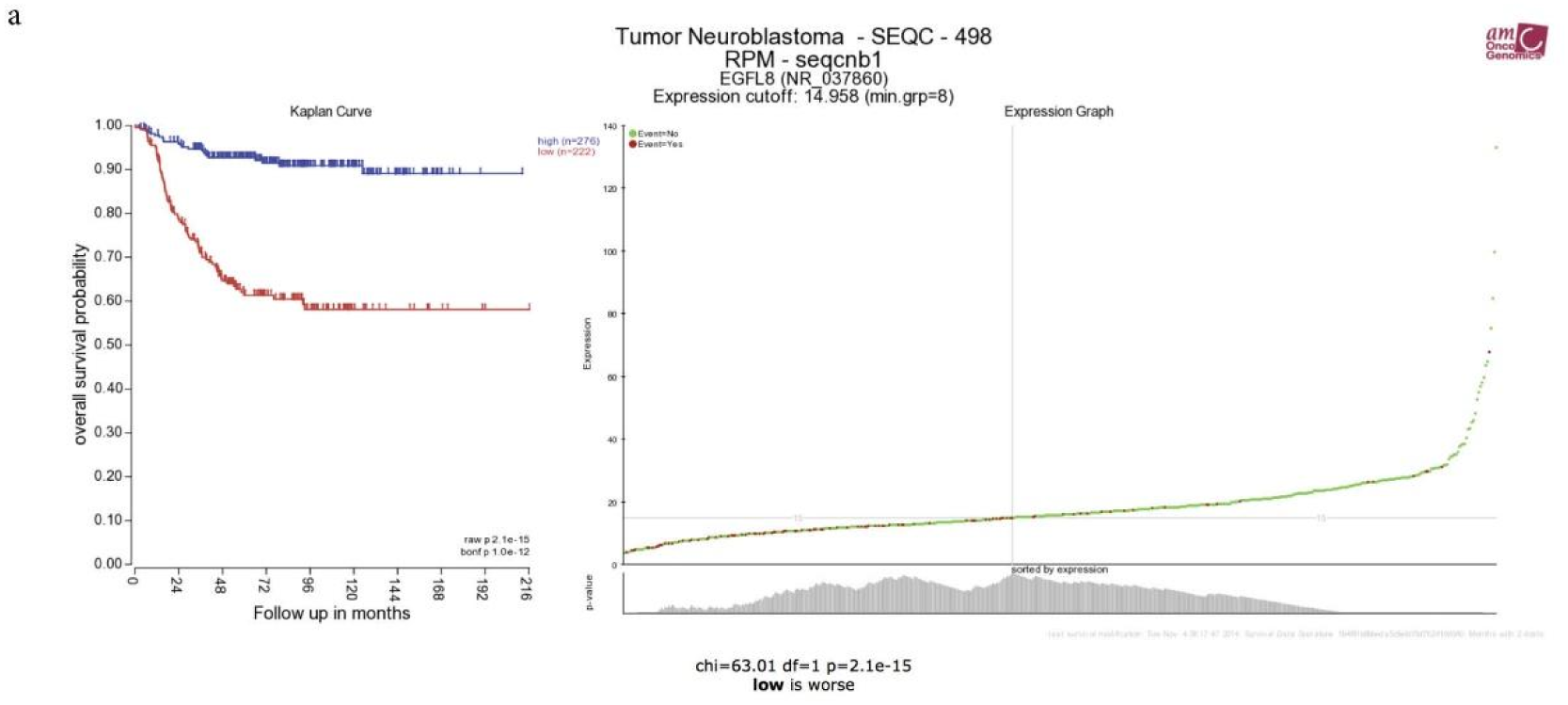
*EGFL8* expression in peripheral neuroblastic tumors. One gene view for *EGFL8* using the *R2 Genomics Analysis and Visualization platform* for the SEQC dataset. **(a)** Kaplan Curve show the overall survival (OS) probability of patients according to *EGFL8* high and *EGFL8* low expressing tumors.

## Notes

### Competing Interest Statement

The authors have declared no competing interest.

